# Human CD4 T cells coordinate antitumor immunity through indirect antigen presentation

**DOI:** 10.64898/2026.09.30.755788

**Authors:** Gargi Roy, Mikel Ruterbusch, Jakob Bakhtiari, Carolyn Shasha, Sylvia M. Lee, Ata S. Moshiri, Connor Wood, Kayla Hopwo, Mikey Witkowski, Julia Szeto, Naina Singhi, Maria Iuliano, Simonne Guenette, Kelly M. McKenna, Evan Newell, Anthony Rongvaux, Joshua R. Veatch

## Abstract

Whether human tumor-specific CD4 T cells can drive antitumor immunity in cancers lacking tumor-cell MHC class II expression has not been established. In MHC class II-negative melanoma and breast cancer, CD4 T cells with a transcriptional signature of tumor specificity correlated with myeloid and CD8 T cell activation. Using a patient-derived NY-ESO-specific, HLA-DP04-restricted TCR, we tested whether human CD4 T cells could control tumors through indirect antigen presentation. In vitro, CD4 T cells inhibited HLA-DP-negative melanoma growth through recognition of tumor-derived antigen on antigen presenting cells (APC). In humanized mice, CD4 T cell transfer induced regression of HLA-DP-negative tumors, and control required HLA-matched human APCs. CD4 T cell transfer induced endogenous CD8 T cell proliferation and myeloid activation that mirrored immune programs correlated with the presence of these cells in patients. These findings establish that human tumor-specific CD4 T cells coordinate antitumor immunity through APC without direct tumor recognition, supporting their therapeutic relevance in MHC class II-negative cancers.

## Introduction

Cytotoxic CD8 T cells recognize peptides presented on class I Major histocompatibility proteins (**MHC I**) which are expressed by nearly all nucleated cells including most cancer cells.^1^ CD4 T cells instead recognize peptide antigens presented on class II MHC (**MHC II**), which is restricted in its expression to professional antigen-presenting cells (APCs) such as dendritic cells (DC), macrophages, and B cells, and is rarely expressed on cancer cells.^2^

Studies in murine models show that CD4 T cells can control tumors through several distinct routes. When tumor cells express MHC class II at baseline or induced by IFN-γ signaling, CD4 T cells can undergo cytolytic differentiation and directly kill tumor cells.^3^ CD4 T cells can also target tumor antigens taken up and presented by DC and other APC to mediate indirect antitumor effects through support of MHC class I restricted tumor specific CD8 T cells^4,5^ and through CD8 T cell independent innate, inflammatory mechanisms.^6–13^ These mechanisms can apply to MHC II negative tumors that do not directly present antigen to CD4 T cells and can control tumors with heterogeneous antigen expression, impaired MHC I antigen presentation, or in contexts where CD8 T cell immunity to tumor antigens is absent.^8,9,13,14^

Murine models differ from human tumors in several important ways that could limit the generalization of murine results to human cancer. Model murine T cell receptors target antigens with higher avidity than clinically relevant targets in human tumors^15,16^ and human and murine CD4 T cells and myeloid cells differ with respect to differentiation plasticity, production of chemokines, and potential myeloid antitumor effector mechanisms such as macrophage production of nitric oxide.^17–19^ Thus, it is not clear which mechanisms identified in murine models apply to human cancers.

Most previous evidence for a role of CD4 T cells in human antitumor immunity has been correlative or anecdotal. We and others previously identified a transcriptional signature of tumor antigen specificity in conventional CD4 T cells in human melanoma^20,21^ marked by expression of the chemokine CXCL13, whose frequency correlated with improved prognosis, greater CD8 T cell proliferation, and elevated myeloid expression of the lymphocyte-recruiting chemokines CXCL9, CXCL10, and CXCL11 and the T cell supporting cytokine IL-15.^20^ Adoptive transfer of tumor-specific CD4 T cells has produced meaningful clinical responses in isolated cases,^22–24^ including a case where transfer of a CD4 T cell product reactive to a tumor neoantigen mediated durable regression of an MHC II negative cholangiocarcinoma.^23^ CD4 T cell reactivity may also play a role in cultured tumor-infiltrating lymphocyte products and neoantigen vaccine responses.^25–27^ These observations implicate CD4 T cells in human antitumor immunity but leave their mechanism, and their causal contribution, undefined.

The majority of work studying the causal therapeutic mechanisms of CD4 T cells in human systems has focused on the ability of CD4 T cells to undergo cytotoxic differentiation and to directly lyse MHC II positive tumor cells in vitro and in vivo in human tumor xenograft models.^28–30^ Whether tumor-specific CD4 T cells targeting clinically relevant cancer antigens can drive antitumor immunity in more prevalent MHC II-negative tumors, where they must rely on indirect presentation, has not been established in a human or humanized system.

Here we show that the presence of conventional CD4 T cells with a transcriptional signature enriched for tumor specificity correlates with activation of myeloid and CD8 T cells in cohorts of melanoma and breast cancer patients without tumor cell expression of MHC II. We isolate a T cell receptor (TCR) from one MHC II negative melanoma patient that recognizes the cancer testes antigen NY-ESO presented by the common HLA-DP04 allele, and demonstrate that TCR transgenic CD4 T cells can mediate control of melanoma through targeting of human APC in an in vitro coculture system and in a humanized mouse model, and that tumor control is dependent on HLA specific presentation of tumor antigen on human APC. A second TCR targeting the neoantigen created by the common BRAF_V600E_ mutation presented by HLA-DQ8 also controlled tumors independently of direct tumor recognition. CD4 T cell transfer activates endogenous CD8 T cells and upregulates expression of CXCL9, CXCL10, CXCL11, and IL15 in myeloid cells, mirroring the transcriptional correlates of tumor antigen specific CD4 T cells in cohorts of melanoma and breast cancer patients. This argues for a causal role of tumor antigen specific non-regulatory CD4 T cells in activating the tumor microenvironment through engagement of APC in human tumors independent of tumor cell presentation to CD4 T cells and supports the therapeutic development of CD4 T cell products and vaccines targeting NY-ESO and other tumor antigens in MHC class II-negative solid tumors to broadly engage antitumor immunity.

## Results

### CXCL13+ CD4 T cells correlate with immune activation in MHC class II-negative tumors

To determine whether tumor antigen-specific CD4 T cells are associated with immune activation in tumors lacking MHC class II expression on cancer cells, we first examined tumor samples from our previously described cohort of 20 melanoma patients (table S1).^20^ Five patients had detectable MHC class II expression on tumor cells by IHC using a pan HLA-DR-DP-DQ antibody, while 15 had minimal or no expression, or expression restricted to infiltrating immune cells, as assessed by a dermatopathologist (Table S1). Single cell transcriptomic analysis revealed that MHC class II expression on tumor cells was associated with a greater fraction of CXCL13+ CD4 T cells and elevated myeloid expression of CXCL9, CXCL10, and CXCL11 as well as a reduced fraction of FOXP3+ regulatory T cells (Fig. 1A–C). However, MHC class II status was not significantly associated with the frequency of proliferating CD8 T cells or overall survival in this small cohort.

**Figure 1.**
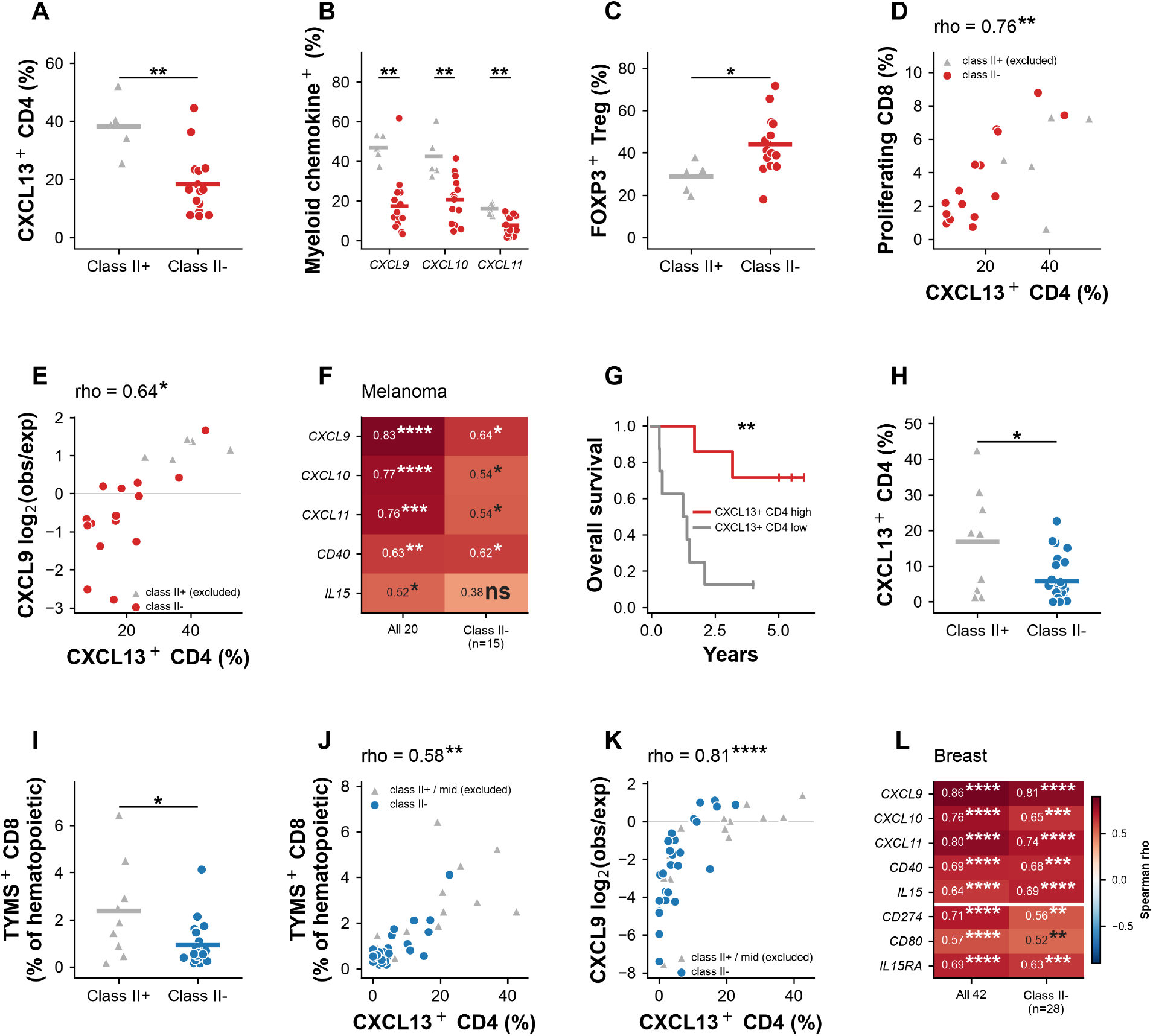
CXCL13+ CD4 T cells, myeloid gene expression and CD8 T cell proliferation in MHC class II-positive and class II-negative tumors. (A–G) Metastatic melanoma cohort (n = 20 patients; MHC class II-positive n = 5, class II-negative n = 15). (A) Fraction of CXCL13+ CD4 T cells, (B) myeloid expression of CXCL9, CXCL10 and CXCL11, and (C) fraction of FOXP3+ regulatory T cells, compared between class II-positive and class II-negative tumors. Horizontal bars are group means. (D) CXCL13+ CD4 T cell frequency versus proliferating CD8 T cell frequency and (E) versus myeloid CXCL9 detection, within class II-negative tumors; excluded class II-positive tumors are shown as grey triangles for context and are not included in the correlation. (F) Correlation of myeloid effector gene detection with CXCL13+ CD4 T cell frequency, in all 20 patients and in the class II-negative subset; values are Spearman ρ and asterisks denote nominal significance (*P < 0.05, **P < 0.01, ***P < 0.001, ****P < 0.0001; ns, not significant). (G) Overall survival of class II-negative patients above (n = 7) and below (n = 8) the median CXCL13+ CD4 T cell fraction; ticks are censored patients. (H–L) Breast cancer cohort (Bassez et al.; n = 42 patients; class II-positive n = 9, intermediate n = 5, class II-negative n = 28). (H) Fraction of CXCL13+ CD4 T cells and (I) frequency of TYMS+ proliferating CD8 T cells, compared between class II-positive and class II-negative tumors. (J) CXCL13+ CD4 T cell frequency versus proliferating CD8 T cell frequency and (K) versus myeloid CXCL9 detection, within class II-negative tumors; excluded class II-positive and intermediate tumors are shown as grey triangles. (L) Correlation of myeloid effector and costimulatory gene detection with CXCL13+ CD4 T cell frequency, in all 42 patients and in the class II-negative subset. In panels e, f, k and l myeloid gene detection is expressed as log2 of the observed detection rate over the rate expected from each cell’s sequencing depth; panel b reports raw percent of cells positive. Group comparisons are by Mann-Whitney U test, correlations are Spearman with permutation P values, and survival is compared by log-rank test. All tests are two-tailed. No correction for multiple comparisons was applied; in (F) and (L) the asterisks denote nominal per-gene significance.

These observations suggest that tumors with MHC II expression on tumor cells associate with a more activated TME, as has been seen previously,^31^ and raise the possibility that the correlation between the presence of CXCL13+ CD4 T cells and activation of myeloid and CD8 T cells could be due to recognition of tumor cells in the subset of cases with MHC II expression.

To ask whether CD4 T cells might coordinate antitumor immunity specifically in MHC class II-negative tumors, we examined correlations between CXCL13+ CD4 T cell frequency and other immune phenotypes within the 15 MHC class II-negative melanoma cases. The fraction of CXCL13+ CD4 T cells correlated with overall survival and myeloid expression of CXCL9, CXCL10, CXCL11, and CD40, with a trend toward correlation with IL15 (Fig. 1D–G) and these genes remained among the most correlated with the fraction of CXCL13+ CD4 T cells (Fig. S1D). These data show that the frequency of CXCL13+ CD4 T cells that are enriched for tumor antigen-specificity are associated with immune activation and favorable outcomes even in the absence of MHC class II expression on cancer cells.

To corroborate these findings in an independent cohort and a second tumor type, we analyzed single-cell RNA sequencing data from 42 breast cancer patients treated with neoadjuvant anti-PD-1 therapy.^32^ Of these, 28 had no little to no detectable MHC class II mRNA expression in tumor cells, 5 had ambiguous expression, and 9 had clear expression. MHC class II-expressing cases showed a higher frequency of CXCL13+ CD4 T cells and TYMS+ proliferating CD8 T cells compared to MHC class II-negative cases, consistent with a more robust immune response when MHC class II was present on tumor cells (Fig. 1H–I); myeloid CXCL10 and CXCL11 expression also differed by tumor MHC class II status (Fig. S1B–C; P = 0.025 and P = 0.041, respectively). Examining only the 28 MHC class II-negative cases, the frequency of CXCL13+ CD4 T cells strongly correlated with myeloid expression of CXCL9, CXCL10, and CXCL11, IL15, CD40, CD274/PD-L1 and CD80 and with the frequency of proliferating (TYMS+) CD8 T cells (ρ = 0.58, P = 0.002) (Fig. 1J–L, Fig. S1E). Taken together, these data suggest that the frequency of tumor antigen-specific conventional CD4 T cells correlates with myeloid cell activation and CD8 T cell proliferation in MHC class II-negative tumors across two independent cohorts and tumor types.

### NY-ESO-specific CD4 T cells infiltrate an MHC class II-negative melanoma tumor

These correlations suggested that tumor-specific CD4 T cells might contribute to immune activation in tumors lacking MHC class II expression on cancer cells. To test whether such cells can drive immune activation and tumor control through indirect antigen presentation, we first defined the antigen specificity and HLA restriction of TCR found in CXCL13+ CD4 T cells from an MHC class II-negative melanoma tumor (Fig. 2A). We had previously identified 8 TCRs specific to 3 different neoantigens created by patient-specific somatic mutations, and 10 TCRs specific to a pool of candidate lineage-specific and cancer-testes antigens known to elicit immune responses in melanoma.^20^

**Figure 2.**
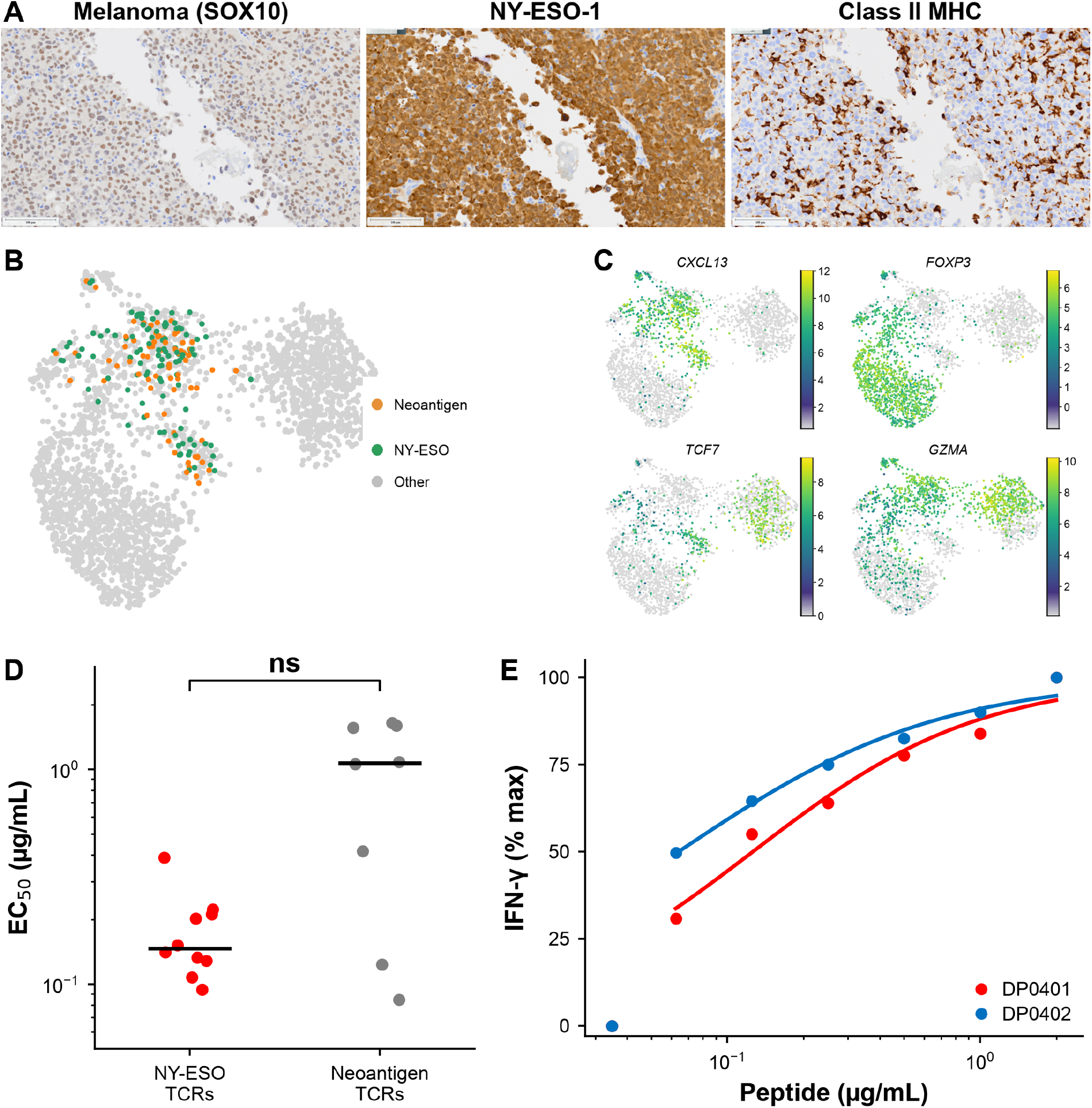
Identification of NY-ESO-1-specific CD4 T cells in MHC class II-negative melanoma. (a) (A) MHC class II expression in the index melanoma patient tumor by immunohistochemistry (SOX10 marks melanoma cells; NY-ESO-1 and MHC class II shown), pathologist-scored as negative on tumor cells and positive on infiltrating myeloid cells. (B) Distribution of NY-ESO-specific and neoantigen-specific CD4 T cells across phenotypic clusters in single-cell RNA sequencing of the primary tumor. (C) Expression of CXCL13, FOXP3, TCF7 and GZMA projected onto the same single-cell RNA sequencing UMAP embedding shown in (B); color indicates normalized expression and is scaled independently for each gene. (D) Functional avidity (EC50) of NY-ESO-specific versus neoantigen-specific TCRs, compared by MWU. (E) IFN-γ dose-response of the lead HLA-DP-restricted TCR (E4) to NY-ESO-1 peptide presented by HLA-DPB1*04:01 (DP0401) and HLA-DPB1*04:02 (DP0402), demonstrating equivalent recognition of both alleles (EC50 DP0401 = 0.13, DP0402 = 0.06 µg/mL). Significance in (D): *P < 0.05, **P < 0.01, ***P < 0.001, ****P < 0.0001; ns, not significant.

To identify the specific shared antigen recognized by the latter group, paired TCRα/β chains were cloned into a lentiviral expression vector and transduced into primary donor CD4 T cells in parallel with CRISPR/Cas9-mediated disruption of the endogenous TCR. Transduced T cells were evaluated for specificity to peptide pools representing each individual antigen by intracellular cytokine staining following coculture with autologous B cells as APCs. All 10 TCRs reacted specifically to a peptide pool from the cancer-testes antigen NY-ESO (Fig. S2A, B). NY-ESO is an attractive immunotherapeutic target as it is shared across patients and tumor types^33^ These NY-ESO-specific TCRs exhibited similar or higher avidity relative to neoantigen-specific TCRs obtained from the same patient (Fig. 2D). Analysis of single-cell RNA sequencing from the primary tumor revealed that NY-ESO-specific and neoantigen-specific CD4 T cells had similar distributions across phenotypic clusters (Fig. 2B). Differential gene expression analysis identified few significant differences, though NY-ESO-specific cells showed modestly higher expression of GZMK and CD27 (Fig. S2D). NY-ESO-specific CD4 T cells thus exhibited avidities and intratumoral differentiation states comparable to those of neoantigen-specific CD4 T cells from the same tumor, supporting their use to investigate the function of naturally occurring human tumor-reactive CD4 T cells.

### A TCR recognizes an NY-ESO epitope presented by either HLA-DP0401 or DP0402

We next determined the HLA restriction of the NY-ESO-1-specific CD4 T cells using B-lymphoblastoid cell lines (B-LCLs) and cell lines expressing HLA alleles shared with the patient. The patient’s HLA class II genotype was HLA-DRB1 01:01/03:01, HLA-DQB1 02:01/05:01, and HLA-DPB1 04:01/04:02. 5 clones reacted to a B-LCL line sharing the DRB1 0301 DQB1 0201 haplotype, and 2 clones reacted to a cell line expressing only HLA-DRB1 0101, and two TCR showed reactivity to both DRB1*01:01 and DQB1*05:01 (Fig. S2C). Notably, one TCR reacted specifically to B-LCL lines sharing only the HLA-DPB1 04:01 or 04:02 alleles and showed similar avidity for both alleles (Fig. 2E). Either HLA-DPB1 4:01 or *04:02 are found in greater than 85% of individuals of European and Hispanic descent and significant fractions of other populations (Table S2),^34^ suggesting this TCR could recognize NY-ESO in a large fraction of patients. An epitope corresponding to NY-ESO residues 161–172 (WITQCFLPVFLA) was identified as critical for TCR recognition (Fig. S2E). This epitope is found in the related CTAG1A, CTAG1B and CTAG2 (LAGE-1a) genes and has been previously described.^35^ Notably, an identical specificity was reported for a CD4 T cell clone that mediated an objective clinical response in a patient with metastatic melanoma.^22^ Alanine scanning demonstrated a W-x-x-Q-C-F-L-P recognition motif that is not found elsewhere in the human proteome, and we did not find evidence for alloreactivity of this TCR across a panel of LCL lines representing different HLA haplotypes. As this TCR could be relevant to the substantial number of patients across tumor types with both NY-ESO expression in tumors and HLA-DP04, we decided to use this to understand mechanisms of human tumor recognition and control by CD4 cells.

### NY-ESO-specific CD4 T cells recognize tumor antigen in vitro through direct and indirect presentation

We asked whether NY-ESO specific CD4 T cells could recognize antigen directly on HLA-DP04+ tumor cells and indirectly though professional antigen presenting cells, and whether these interactions could mediate tumor control. TCR-transgenic CD4 T cells produced IFN-γ when cocultured with the melanoma cell line SK-MEL-37 that endogenously expresses NY-ESO and HLA-DP*04:01 SK-MEL-37 but did not recognize SK-MEL-37 cells following CRISPR-mediated disruption of either NY-ESO or HLA-DP (Fig. 3A). Furthermore, TCR-transgenic cells mediated killing of SK-MEL-37 cells in a manner dependent on target cell expression of HLA-DP (Fig. 3B). These results demonstrate that CD4 T cells expressing this TCR can directly recognize and kill melanoma cells presenting endogenous levels of NY-ESO and HLA-DP.

**Figure 3.**
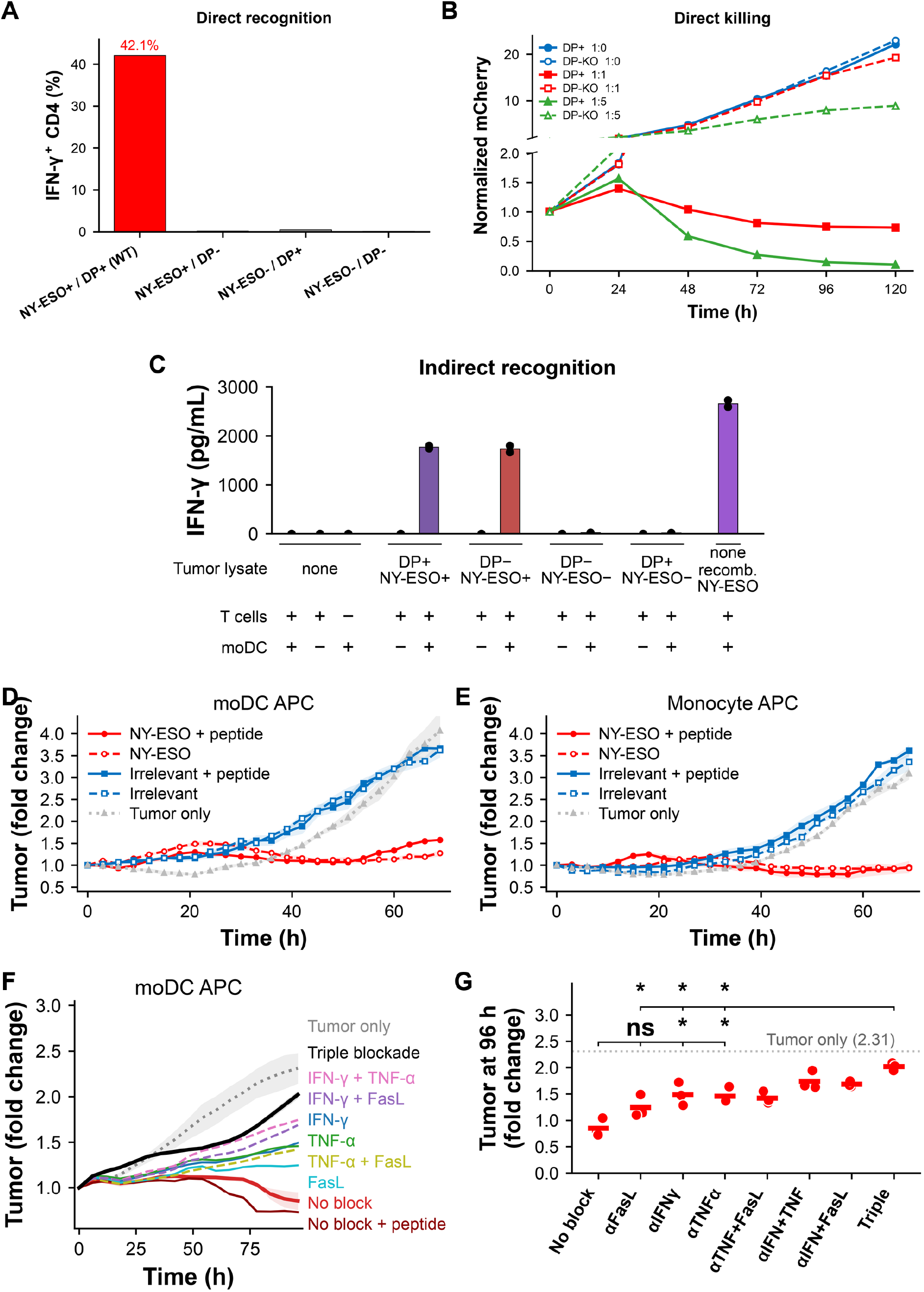
NY-ESO-1-specific CD4 T cells recognize tumor antigen through direct and indirect presentation. (a) (A) Direct recognition of SK-MEL-37 melanoma variants by NY-ESO-specific TCR-transduced CD4 T cells, measured as the percentage of IFN-γ⁺ CD4 T cells after coculture. Recognition of wild-type (NY-ESO-1⁺/HLA-DP⁺) tumor cells (42.1%) was lost upon CRISPR knockout of either NY-ESO or HLA-DP, or both (<0.5%). EGFR-and Merkel-specific control TCRs were unreactive in the same experiment (not shown). Direct recognition requires both cognate antigen and HLA-DP on the tumor cell. (B) Growth of SK-MEL-37 HLA-DP-positive or HLA-DP knockout targets cocultured with NY-ESO-specific CD4 T cells at effector:target ratios of 0:1, 1:1 and 5:1, by IncuCyte live-cell imaging (Red Object Integrated Intensity, RCU × μm²/well, fold change from t = 0). (C) Indirect recognition: IFN-γ secretion by NY-ESO-specific CD4 T cells cocultured with HLA-DPB1*04:01⁺ monocyte-derived dendritic cells (moDCs) pulsed with lysate from SK-MEL-37 variants. (D, E) Indirect control of mCherry⁺ SK-MEL-37 HLA-DP knockout tumor cells co-cultured with NY-ESO-specific or irrelevant-TCR CD4 T cells, with or without exogenous NY-ESO peptide, versus tumor-only control, using (D) moDCs or (E) peripheral blood monocytes as APCs (IncuCyte, fold change from 0 h, over 70 h). (F) Time course of the moDC indirect control assay under blockade of IFN-γ, TNF-α and FasL, alone and in combination, with no-block and tumor-only references, over 96 h. (G) Endpoint (96 h) tumor fold change across single and combined blockade with moDC APCs; dotted line = moDC tumor-only reference; horizontal bars are arithmetic means. n = 3 wells per condition in (D–G), except the moDC tumor-only reference (n = 2). In (G), endpoint tumor fold change was compared between conditions by two-tailed Welch’s t test on n = 3 replicate wells per condition without correction for multiple comparisons. (*P < 0.05, **P < 0.01, ***P < 0.001, ****P < 0.0001; ns, not significant).

As the patient from whom this TCR was identified did not express MHC class II on tumor cells, we hypothesized that antigen recognition in vivo occurred through professional APCs that acquire and process tumor-derived material. To test this, we cocultured TCR-transgenic CD4 T cells with HLA-DP04:01-expressing monocyte-derived dendritic cells (moDCs) that had been pulsed with crude lysate from SK-MEL-37 variants. IFN-γ production was dependent on NY-ESO expression in the tumor cells from which the lysate was derived but was independent of HLA-DP expression in the lysate source cells (Fig. 3C). This demonstrates that the TCR can mediate indirect antigen recognition through professional APCs.

Although lysate-pulsed moDCs established that NY-ESO could be processed and presented indirectly, this experiment did not determine whether APCs could acquire sufficient antigen directly from tumor cells to activate CD4 T cells or whether such recognition could mediate tumor control. To address this, we established a coculture containing mCherry-expressing HLA-DP-knockout SK-MEL-37 cells, NY-ESO-specific CD4 T cells, and HLA-DP04-positive peripheral blood monocytes or monocyte derived dendritic cells (moDC) and monitored tumor growth by live-cell imaging. In the presence of either APC population, NY-ESO-specific CD4 T cells inhibited the growth of HLA-DP-knockout tumor cells, whereas tumor cells grew progressively in control conditions with an irrelevant TCR or lacking APC (Fig. 3D–E). This occurred independently of exogenous peptide antigen. Thus, monocytes and moDCs acquired and presented sufficient endogenously expressed NY-ESO from tumor cells to initiate tumor control. Because the tumor cells could not present antigen to the CD4 T cells in the coculture and CD8 T cells and NK cells were absent, these findings establish that indirect antigen presentation is sufficient to control tumor growth without direct recognition of cancer cells.

We next asked which effector mechanisms downstream of indirect antigen recognition might control tumor growth. Blockade of IFN-γ, TNF-α, or FasL individually produced partial restoration of tumor growth, whereas simultaneous blockade of all three pathways restored tumor growth to approximately that observed in tumor-only cultures (Fig. 3F–G). Similar effects were observed with monocytes and moDCs as APCs (Fig. S3A, B). These data indicate that indirect recognition initiates partially overlapping IFN-γ-, TNF-α-, and FasL-dependent mechanisms.

### NY-gESO-specific CD4 T cells mediate APC-dependent control of MHC class II-negative tumors in humanized mice

While indirect antigen presentation by APCs has been shown to mediate CD4 T cell-dependent tumor control in conventional mouse models,^4,7,8,36,37^ standard human tumor xenograft systems lack human APCs, precluding dissection of these mechanisms in the context of a human immune system. To address this, we utilized the MISTRG humanized mouse model, which carries knock-ins for human M-CSF, IL-3/GM-CSF and TPO together with a human SIRPA transgene, permitting engraftment and multilineage differentiation of human hematopoietic stem cells (HSCs) into human T, B, and myeloid cells.^38^ Newborn MISTRG mice engrafted with HLA-DP04-positive fetal liver CD34+ HSCs and implanted subcutaneously with SK-MEL-37 melanoma form tumors infiltrated with human T cells, dendritic cells (DCs), and tumor-associated macrophages (TAMs) (Fig. 4A). This system therefore provides a tractable platform in which human CD4 T cells can interact with human APCs in the context of a human tumor microenvironment.

**Figure 4.**
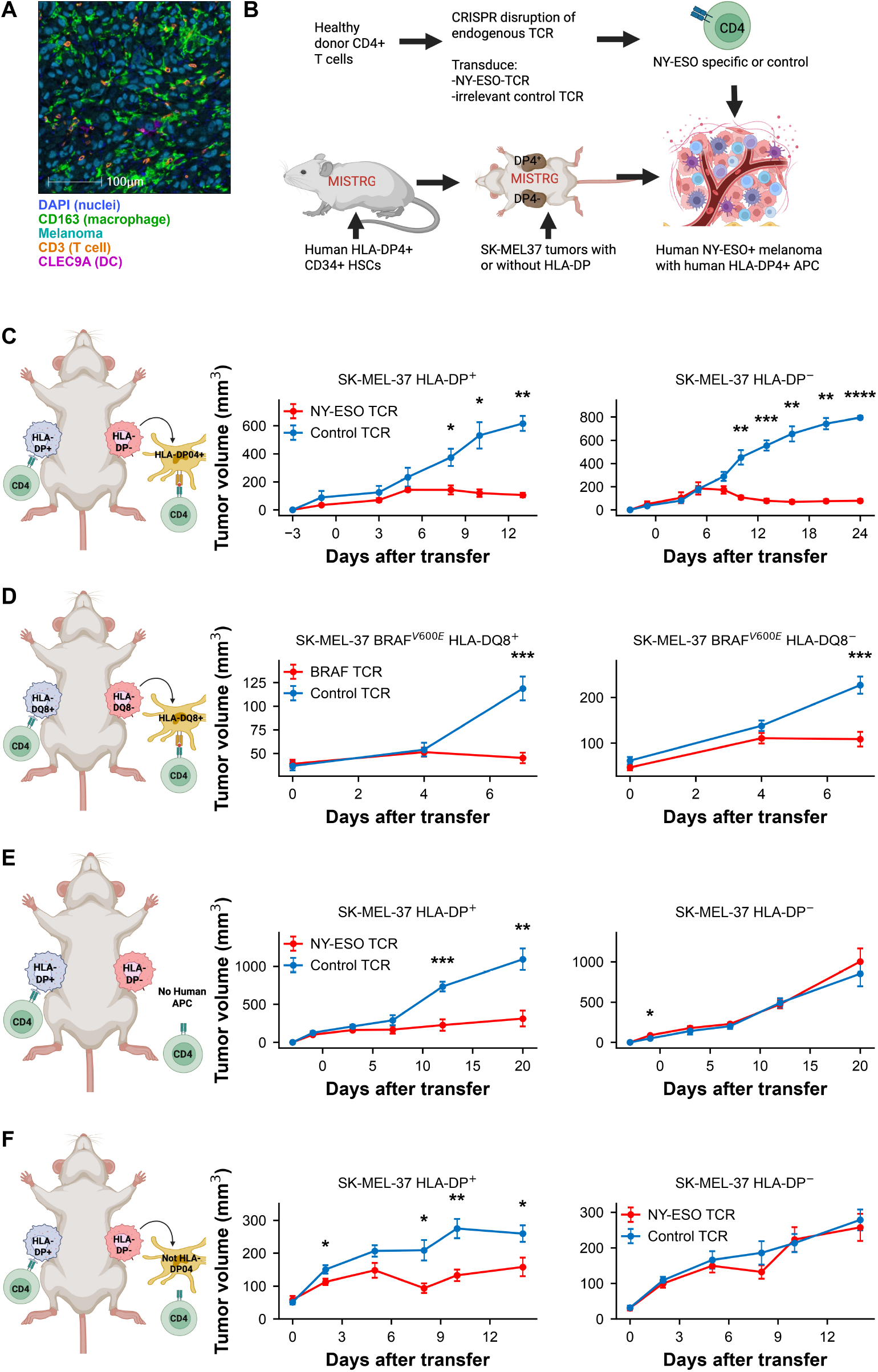
Tumor control by NY-ESO-1-specific and BRAF_V600E_-specific CD4 T cells in humanized MISTRG mice with and without matched human APCs. (A) Multiplex immunofluorescence of an SK-MEL-37 tumor from an HLA-DP04-positive humanized MISTRG mouse, showing human macrophages (CD163), T cells (CD3), dendritic cells (CLEC9A), melanoma cells, and nuclei (DAPI). (B) Schematic of the experimental design. (C) Tumor volume over time in MISTRG mice engrafted with HLA-DP04-positive HSCs and bearing bilateral SK-MEL-37 tumors (HLA-DP-positive and HLA-DP knockout flanks of the same animal) following adoptive transfer of NY-ESO-specific or irrelevant control TCR CD4 T cells (n = 5 NY-ESO TCR and 6 control TCR mice). (D) Tumor volume over time in MISTRG mice engrafted with HLA-DQ8-positive HSCs and bearing bilateral SK-MEL-37 tumors over-expressing the BRAF_V600E_ mutation minigene fused to GFP (HLA-DQ8-transduced and HLA-DQ8 negative flanks), following adoptive transfer of CD4 T cells expressing a BRAF_V600E_ HLA-DQ8-restricted TCR or control TCR (n = 9 BRAF TCR and 7 control TCR mice). (E) Tumor volume in non-humanized MISTRG mice (lacking human APCs), bearing bilateral SK-MEL-37 HLA-DP-positive and HLA-DP knockout tumors treated with NY-ESO-specific CD4 T cells (n = 7 mice per arm). (F) Tumor volume in MISTRG mice engrafted with HLA-DP04-negative (mismatched) HSCs, bearing bilateral SK-MEL-37 HLA-DP-positive and HLA-DP knockout tumors treated with NY-ESO-specific CD4 T cells (n = 9 NY-ESO TCR and 12 control TCR mice). Data are mean ± SEM; curves are shown through the last day on which both arms retained at least three mice (individual-mouse curves and later timepoints in Fig. S4A, D–F). Control (blue); NY-ESO-specific or BRAF_V600E_-specific (red). In c–f, asterisks denote nominal significance for the difference in tumor volume between arms at that timepoint by Welch’s t test, with no correction for multiple comparisons (*P < 0.05, **P < 0.01, ***P < 0.001, ****P < 0.0001; ns, not significant).

To test whether NY-ESO-specific CD4 T cells can control tumors and its dependence on HLA-DP04 expression on tumor cells, we engrafted MISTRG mice with HSCs from an HLA-DP04-positive donor and implanted parental SK-MEL-37 tumors (capable of direct antigen presentation to NY-ESO-specific CD4 T cells) in one flank and SK-MEL-37 HLA-DP knockout tumors generated by CRISPR/Cas9 (incapable of direct presentation) in the contralateral flank of the same animal. Adoptive transfer of NY-ESO-specific but not control CD4 T cells mediated regression of both HLA-DP-positive and HLA-DP knockout tumors (Fig. 4C), demonstrating that NY-ESO-specific CD4 T cells can control tumors that are unable to directly present cognate antigen, although 2 mice showed delayed outgrowth of DP-tumors (Fig. S4A), and depth as well as durability of control varied with HSC and CD4 T cell donor (Fig. S4B, C). To test whether indirect tumor control is a general property of tumor-specific CD4 T cells rather than a peculiarity of the NY-ESO/HLA-DP04 pairing, we independently confirmed these findings in a second, non-overlapping HLA class II system. We used a TCR specific for the BRAF_V600E_ neoantigen presented by HLA-DQ8^16^ and MISTRG mice engrafted with HLA-DQ8-positive HSCs bearing bilateral mutant SK-MEL-37 tumors expressing the BRAF_V600E_ antigen linked to GFP. As with the NY-ESO system, adoptive transfer of BRAF_V600E_-specific CD4 T cells controlled both HLA-DQ8-positive tumors (capable of direct presentation) and HLA-DQ8 negative tumors (incapable of direct presentation) (Fig. 4D). That a distinct tumor antigen, restricted by a distinct HLA class II allele, reproduces control of tumors unable to present antigen directly establishes APC-dependent indirect control occurs in this humanized model with multiple clinically relevant shared tumor antigens.

To determine whether tumor control of HLA-DP-negative tumors required engagement of human APCs through the same HLA-DP04 restriction, we repeated the NY-ESO TCR transfer in non-humanized MISTRG mice that lack human APCs, and in MISTRG mice engrafted with HSC that do not express HLA-DP04. In both settings, NY-ESO-specific CD4 T cells controlled parental SK-MEL-37 tumors capable of direct antigen presentation, although less efficiently than in HLA-DP04-positive humanized mice, but failed to control HLA-DP knockout tumors (Fig. 4E, F). Taken together, these data demonstrate that NY-ESO-specific CD4 T cells mediate tumor control of MHC class II-negative tumors through engagement of human APCs presenting the cognate antigen on the matched HLA class II allele. Individual-mouse tumor trajectories and the full observation period for all arms, including timepoints beyond those shown in Fig. 4, are provided in Fig. S4A, D–F.

### NY-ESO specific CD4 T cells are locally activated in HLA-DP+ and HLA-DP-negative tumors and activate endogenous CD8 T cells

Having shown that tumor-specific CD4 T cells can control tumors through antigen recognition on human APCs, we asked whether their transfer induces the CD8 and myeloid activation associated with CXCL13-positive CD4 T cells in patients. To enable tracking of transferred cells, NY-ESO-specific and control TCR constructs were co-transduced with an epitope tag, and tumors were harvested 8 days after T cell transfer for analysis of transferred and endogenous human immune populations by flow cytometry. There was variability in the frequency of human myeloid cells, CD8+ T cells, regulatory T cells, and transferred T cells in tumors across our cohort (Fig. S5A-C).

To restrict comparisons of transferred CD4 T cell phenotype to tumors with adequate engraftment of transferred cells, we excluded samples in which transferred (NGFR+) cells comprised less than 2% of the total CD4 T cell compartment. Transferred NY-ESO specific and control CD4 T cells showed little to no expression of FoxP3, arguing against regulatory differentiation of the transferred cells (Fig. S5D). Relative to control CD4 T cells transduced with an irrelevant TCR, transferred NY-ESO-specific CD4 T cells showed increased expression of markers of antigen engagement (PD-1, CTLA-4), proliferation (KI-67), activation (HLA-DR), and effector function (GZMB) relative to control TCR-transduced cells in both HLA-DP-positive and HLA-DP knockout flanks of the same animals (Fig. 5A), with the magnitude of induction broadly comparable across the two tumor types. These data indicate that transferred CD4 T cells are activated locally in both HLA-DP-positive and HLA-DP knockout tumors, and that engagement of cognate peptide-MHC complexes on human APCs is sufficient to drive a functional CD4 T cell response in the absence of direct presentation by tumor cells.

**Figure 5.**
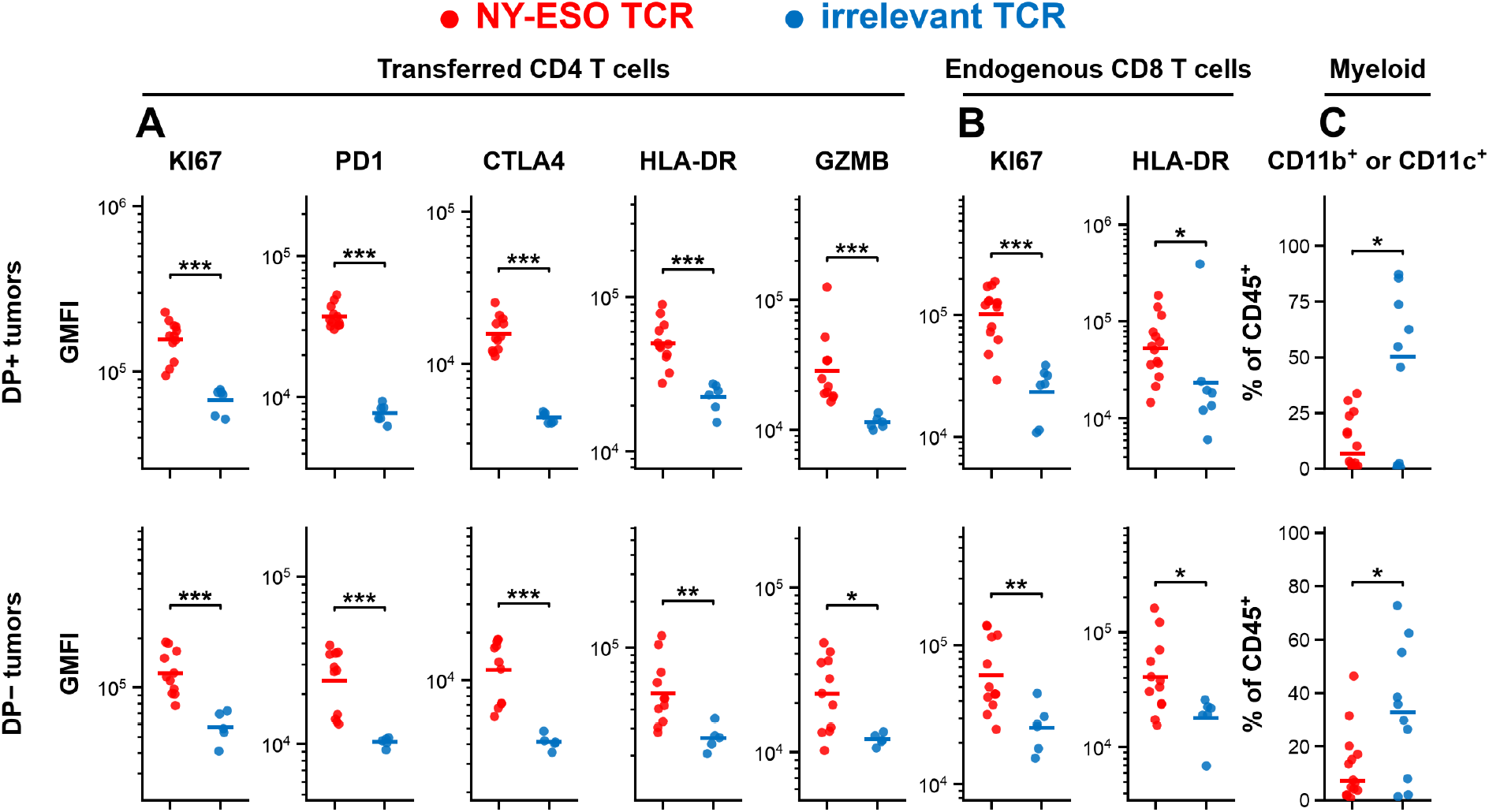
NY-ESO-specific CD4 T cells are activated locally in tumors independent of direct antigen recognition on tumor cells. Flow cytometric analysis of tumor-infiltrating immune cells 8 days after adoptive transfer of NGFR-tagged NY-ESO-specific or control TCR CD4 T cells into HLA-DP04-positive humanized MISTRG mice bearing bilateral HLA-DP-positive (top row) and HLA-DP knockout (bottom row) SK-MEL-37 tumors. (A) Geometric mean fluorescence intensity (GMFI) of KI-67, PD-1, CTLA-4, HLA-DR, and GZMB on transferred (NGFR+) CD4 T cells; only tumors in which transferred cells comprised at least 2% of the total CD4 T cell compartment were included (n = 12 NY-ESO DP+, 11 NY-ESO DP−, 6 Control DP+, 5 Control DP−). (B) GMFI of KI-67 and HLA-DR on endogenous CD8 T cells; (n = 14 NY-ESO DP+, 12 NY-ESO DP−, 7 Control DP+, 6 Control DP−). (C) Frequency of CD3− CD11b+ or CD11c+ myeloid cells as a fraction of CD45+ cells (n = 14 NY-ESO DP+, 14 NY-ESO DP−, 10 Control DP+, 10 Control DP−). Filled red circles indicate NY-ESO-specific CD4 T cell-treated tumors; filled blue circles indicate control TCR-treated tumors. Horizontal bars indicate the geometric mean in (A) and (B) and the median in (C). Significance (*P < 0.05, **P < 0.01, ***P < 0.001, ****P < 0.0001; ns, not significant) is from two-tailed exact Mann-Whitney U tests for the GMFI panels in (A) and (B), and from two-tailed Welch’s t test for the frequency panels in (C); not corrected for multiple comparisons.

To determine whether transferred CD4 T cells provide help to endogenous CD8 T cells in this system, we examined endogenous human CD8 T cells in the same tumors. Endogenous CD8 T cells from NY-ESO TCR-treated mice showed increased expression of KI-67 and HLA-DR relative to controls in both HLA-DP-positive and HLA-DP knockout flanks (Fig. 5B), consistent with CD4-mediated help driving CD8 T cell proliferation and activation in the tumor. The induction of CD8 T cell proliferation in HLA-DP knockout tumors, where direct presentation of NY-ESO to CD4 T cells by tumor cells is precluded, indicates that CD4 T cell help to endogenous CD8 T cells is delivered indirectly through APCs.

To examine whether CD4 T cell transfer also affects the tumor myeloid compartment, we measured the frequency of CD3− CD11b+ or CD11c+ myeloid cells as a fraction of CD45+ cells in the same tumors by flow cytometry. NY-ESO TCR transfer was accompanied by a reduction in the frequency of CD3− CD11b+ or CD11c+ myeloid cells in both HLA-DP-positive and HLA-DP knockout tumors (Fig. 5C), which could be downstream of either depletion of myeloid populations or increased recruitment of lymphoid populations.

### NY-ESO specific CD4 T cells have a distinct non-regulatory transcriptional phenotype in HLA-DP negative tumors

To extend the flow cytometric findings and define the immune-cell states associated with CD4 T cell therapy, we performed targeted single-cell RNA sequencing with paired TCR sequencing on sorted human CD45+ cells from HLA-DP-knockout tumors 8 days after transfer of NY-ESO-specific or irrelevant EGFR-specific CD4 T cells into HLA-DP04-positive MISTRG mice (n = 6 mice per group; 17,088 cells called, of which 14,204 carried an assigned sample tag and 13,379 were retained in the clusters shown in Fig. 6A; 404-gene targeted immune-response and 30-marker AbSeq panels). Dimensional reductions with UMAP demonstrated a CD8 T cell population, a FOXP3+ IL2RA+ regulatory T cell population, and three conventional CD4 T cell states (Fig. 6A, B). PD-1-high conventional CD4 T cells showed expression of CXCL13 separated into TCF7-high and TCF7-low populations, whereas a third population expressed lower PD-1. Myeloid and rare B-lineage populations were also identified.

**Figure 6.**
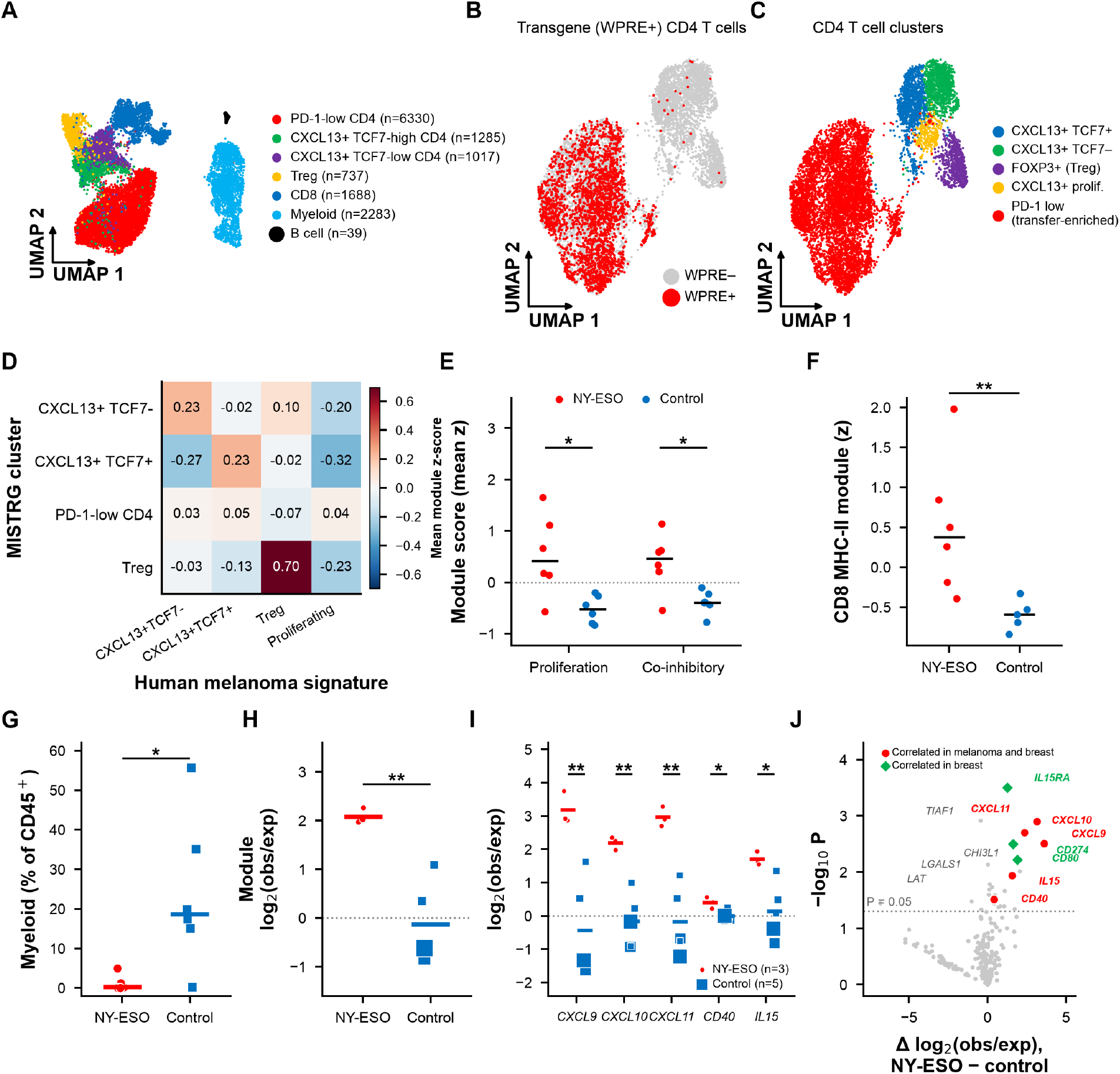
Single-cell RNA sequencing of NY-ESO TCR-treated tumors recapitulates the flow cytometric findings and identifies CD4-driven myeloid chemokine induction. Targeted scRNA-seq with paired TCR sequencing of FACS-sorted human CD45+ cells from SK-MEL-37 HLA-DP knockout tumors 8 days after adoptive transfer of NY-ESO-specific or irrelevant EGFR-specific control TCR CD4 T cells into HLA-DP04-positive humanized MISTRG mice (n = 6 mice per group). (A) UMAP projection of 13,379 human CD45+ cells, colored by cluster annotation. (B) UMAP overlay of WPRE-positive transferred cells. (C) UMAP of the CD4 T cell compartment colored by subset. (D) Module scores for human melanoma CD4 subset signatures (top 30 genes per signature) across MISTRG CD4 T cell clusters. (E) Proliferation (PCNA, MCM2, MCM4, TYMS, MKI67, TOP2A and PTTG2) and co-inhibitory receptor module (CTLA4, PDCD1, LAG3, HAVCR2, TIGIT, BTLA, CD160, CD244, ENTPD1 and TOX) scores in WPRE-positive cells within the PD-1 low CD4 cluster, NY-ESO versus control TCR-treated tumors (n = 6 mice per group). Each mouse contributes its mean log(1+count) per gene across its WPRE-positive cells, for mice with at least two such cells; each gene is standardized across the twelve mice and the module score is the mean of the standardized values over the module’s genes. Points are mice, horizontal bars are group medians. (F) Composite MHC class II module score (HLA-DRA + HLA-DPA1 + HLA-DQB1 + HLA-DMA + CD74) on CD8 T cells, NY-ESO versus control TCR-treated tumors (NY-ESO n = 6, control n = 5 mice; one control mouse yielded fewer than two CD8 T cells and was excluded). (G) Frequency of myeloid cells as a fraction of CD45+ cells in the scRNA-seq dataset, NY-ESO versus control TCR-treated tumors (n = 6 mice per group); horizontal bars are group medians. (H) Composite myeloid effector module (CXCL9 + CXCL10 + CXCL11 + CD40 + IL15) and (I) the individual genes, expressed as detection over depth-based expectation in tumor myeloid cells for mice with at least 5 myeloid cells (NY-ESO n = 3, control n = 5 mice. (J) Differential detection corrected for cell sequencing depth across all 220 measurable genes in tumor myeloid cells (NY-ESO n = 3, control n = 5 mice). Significance marks (*P < 0.05, **P < 0.01, ***P < 0.001, ****P < 0.0001; ns, not significant), uncorrected for multiple comparisons, are from Welch’s t test on per-mouse values, except E, F and G, which used two-tailed exact Mann-Whitney U, and LAT in J, for which all three NY-ESO mice were at zero and the value shown is a two-tailed exact Mann-Whitney U on the per-mouse detection fractions (P = 0.036, the smallest value attainable with three versus five mice).

The CD4 T cell states resembled those previously identified in human melanoma. The TCF7-high and TCF7-low PD-1-high populations shared features with the progenitor-like and more differentiated CXCL13-positive CD4 T cells, whereas the regulatory T cell cluster expressed the expected FOXP3, IL2RA, and CTLA4 program; module scores built from the corresponding human melanoma CD4 signatures^20^ separated the MISTRG clusters along the same axes (Fig. 6D). The presence of these recognizable populations is notable given important differences between this model and human tumors, including potential xenogeneic or alloreactive T cell responses, absent human lymphoid architecture, and abnormal thymic selection.^39^

Cells expressing the WPRE element from the TCR transgene were enriched more than 20-fold in the PD-1 low CD4 T cell cluster compared with the PD-1 high CD4, CD8, regulatory T cell, and myeloid populations (38.1% versus 0.3–1.0%; Fig. 6B). Due to limitations in detection in single cells, this value provides a lower bound on the transferred-cell content of the cluster and implies that transferred CD4 T cells could make up a large fraction of cells in this cluster, and few or no cells in other clusters. The PD-1 low CD4 population differed from the endogenous PD-1 high conventional CD4 T cell cluster with lower expression of genes related to chronic antigen stimulation and higher expression of genes related to cytotoxic differentiation (TOX, LAG3, GZMA and GZMB shown in Fig. S6A). This indicated that transferred cells take on a distinct and potentially less exhausted phenotype from endogenous conventional CD4 T cells in this model, and do not overlap in phenotype with regulatory T cells.

Comparison of WPRE+ cells within the PD-1 low CD4 cluster between tumors treated with the NY-ESO TCR and tumors treated with the control TCR showed increased expression of proliferation associated genes as well as coinhibitory receptor genes (Fig. 6E), consistent with the antigen recognition and activation measured by flow cytometry (Fig. 5A), demonstrating that NY-ESO specific CD4 T cells were locally activated in HLA-DP04 negative tumors.

### The myeloid transcriptional program associated with CXCL13+ CD4 T cells in human cancer is induced by NY-ESO specific CD4 T cells in HLA-DP negative tumors

Endogenous CD8 T cells from NY-ESO-treated tumors also showed evidence of increased activation. A composite score of MHC class II genes (HLA-DRA, HLA-DPA1, HLA-DQB1, HLA-DMA and CD74) was higher in NY-ESO relative to control TCR-treated tumors (Fig. 6F; individual module components per mouse in Fig. S6B). This finding was consistent with increased Ki-67 and HLA-DR expression on endogenous CD8 T cells by flow cytometry (Fig. 5B) and suggests enhanced proliferation of CD8 T cells following NY-ESO specific CD4 T cell transfer.

NY-ESO TCR transfer was also associated with a lower frequency of human myeloid cells among captured CD45+ cells (Fig. 6G). This result was consistent with the reduction in CD3-negative CD11b+ or CD11c+ cells measured by flow cytometry (Fig. 5C) but cannot distinguish loss of myeloid cells from increased relative representation of T cells. Myeloid expression of CXCL9, CXCL10, CXCL11, CD40 and IL15 correlated with the frequency of CXCL13-positive CD4 T cells in human melanoma and breast cancer,^20^ (Fig. 1), so we hypothesized that transfer of NY-ESO specific CD4 T cells would increase expression of these genes in myeloid cells. Among tumors with sufficient myeloid cells for mouse-level analysis (3 of 6 NY-ESO-treated and 5 of 6 control tumors, the retained NY-ESO tumors contributing 6, 18 and 29 myeloid cells), NY-ESO treatment increased the sequencing depth-adjusted fraction of myeloid cells expressing each of these five genes (Fig. 6H-J). IL15RA, CD274 (PD-L1) and CD80 were also nominally increased, and correlated with the presence of CXCL13+ CD4+ T cells in the breast cancer but not melanoma cohorts (Fig. 1, Fig. S1D-E). Concordant induction of this human myeloid signature links antigen-specific CD4 T cell transfer to the inflammatory program associated with CXCL13-positive CD4 T cells in MHC class II-negative human tumors.^20^ Together, the single-cell data were concordant with the flow cytometric changes in transferred CD4, endogenous CD8, and myeloid compartments and support a model where tumor specific CD4 T cells play a causal role in activation of myeloid and CD8 immunity in human melanoma that is independent of direct recognition of antigen on tumor cells.

## Discussion

The contribution of tumor antigen specific CD4^+^ T cells to human antitumor immunity remains poorly understood, particularly in tumors that lack class II MHC expression on cancer cells. Murine studies have established that CD4^+^ T cells can control such tumors through recognition of antigen on APCs,^6–8,37,40^ but standard human tumor xenograft models do not contain the human myeloid populations needed to test these interactions.^38,39^ The data presented here show that human CD4^+^ T cells recognizing a naturally expressed tumor antigen can mediate regression of tumors that cannot directly present the antigen through engagement of HLA-matched human APCs. Our finding that CD4^+^ T cell transfer also induced myeloid and CD8^+^ T cell activation similar to patterns associated with tumor reactive CD4^+^ T cells in patients^20^ argues for a role of these cells in coordinating human antitumor immunity.

CD4^+^ T cells can support antitumor immunity in murine models through several mechanisms that depend on antigen presentation by APCs. CD4^+^ T cells support the function of tumor specific CD8^+^ T cells,^4^ including through interactions with dendritic cells within the tumor,^5,41^ and that activation of myeloid cells by CD4^+^ T cells can mediate tumor control through inflammatory effects on cancer cells and the tumor vasculature.^8–10,12,13,36^ These findings suggest that targeting tumor antigens with CD4^+^ T cells could augment CD8^+^ T cell responses and provide additional antitumor effects when direct recognition of cancer cells is impaired of endogenous CD8 T cell responses are absent.^8,13^ However, human and murine myeloid cells differ in their responses to inflammatory signals. For example, nitric oxide is a critical contributor to inflammatory tumor control in murine models,^8,10^ whereas human macrophages produce substantially less nitric oxide in response to inflammatory stimulation.^18^ Also, the functional avidity of model murine TCR and antigens may not be representative of clinically relevant human antigen targets.^15,16,42^ Thus, the ability of human CD4^+^ T cells to recognize tumor antigen on APCs does not by itself establish that the effector mechanisms identified in mice will be effective in human tumors. Our data showing APC-dependent control with human CD4^+^ T cells and human myeloid cells provide a basis for testing which of these mechanisms contribute to human antitumor activity. Our data suggest that either monocyte or monocyte derived dendritic cells can take up and present endogenous levels of antigen in vitro to activate CD4 T cells, and mediate tumor control through additive effects of FASL, IFN-γ and TNF-α, and this system given an opportunity to further dissect inflammatory mechanisms of tumor control.

We previously found that the frequency of conventional CD4^+^ T cells with a signature of tumor antigen specificity correlated with CD8^+^ T cell proliferation, an immune stimulatory phenotype in myeloid cells, and improved survival in human melanoma.^20^ These observations suggested diverse functional roles for tumor specific CD4^+^ T cells but did not establish whether they were responsible for the associated changes in the tumor microenvironment. Our finding that the associations with myeloid and CD8^+^ T cell activation persist in melanoma and breast cancers without class II MHC on tumor cells supports a role for CD4^+^ T cells beyond direct recognition of cancer cells. Furthermore, the induction of myeloid CXCL9, CXCL10, CXCL11, CD40, and IL15 and activation of endogenous CD8^+^ T cells following CD4^+^ T cell transfer provides experimental support for the human CD4 T cells driving changes in the human melanoma TME rather than being a consequence.

Our study is limited by variability in human immune engraftment in MISTRG mice,^38^ incomplete HLA matching between the hematopoietic donor and tumor, and development of endogenous human T cells in a mouse thymus. Although endogenous CD8^+^ T cells were activated following CD4^+^ T cell transfer, their antigen specificity and contribution to tumor control remain undefined, and we cannot exclude contributions from alloreactive or xenogeneic responses. The model also does not reproduce the human stromal and vascular compartments, which could influence the effects of inflammatory signals on tumor growth. Nevertheless, the presence of human macrophages and dendritic cells in MISTRG tumors allowed us to test the requirement for presentation of tumor antigen on HLA-matched human APCs, which is not possible in standard human tumor xenografts. The ability to manipulate antigen presentation in the tumor and hematopoietic compartments provides an approach to determine which human APC populations support CD4^+^ T cell activity and how myeloid and endogenous T cell responses contribute to tumor control.

Immunotherapies that augment CD4^+^ T cell responses to shared tumor antigens are attractive because they could be applied to multiple patients and engage several components of antitumor immunity. NY-ESO is a clinically relevant target for this approach, as it is expressed in multiple tumor types,^33^ can be acquired and presented by human APCs, and has been associated with a clinical response following transfer of NY-ESO specific CD4^+^ T cells.^22^ Recognition of antigen presented by HLA-DP0401 or 0402 that are collectively present in over 85% of Americans of either European or Hispanic ancestry (Table S2)^34^ will make clinical trials targeting NY-ESO with CD4 T cells more feasible, and we are in the process of starting a phase I clinical trial of adoptive transfer of CD4 T cells in NY-ESO expressing cancer. Our observation of indirect tumor control with a second TCR recognizing the BRAF_V600E_ that is present in 40% of advanced melanoma patients through HLA-DQ8^16^ provides additional support for targeting APC-presented tumor antigens. This work establishes the therapeutic relevance of CD4 T cells in activation of the tumor microenvironment in human cancer independently of tumor cell class II MHC expression and suggests the possibility of therapies targeting class II MHC antigens.

## Methods

### Patient samples and single-cell RNA sequencing analysis

Melanoma patient samples and single-cell RNA sequencing data were obtained from our previously published cohort.^20^ MHC class II expression on tumor cells was determined by review of archival tumor sections stained with HLA-DR-DQ-DP antibodies by IHC and reviewed by a dermatopathologist. Classification of patients as MHC class II-positive (n = 5) or MHC class II-negative/low (n = 15) was based on the absence of strong staining on malignant cells; the negative/low group includes cases with weak, heterogeneous staining largely restricted to admixed immune cells. Breast cancer single-cell RNA sequencing data were obtained from Bassez et al., comprising 42 patients treated with neoadjuvant anti-PD-1 therapy. MHC class II status was classified from a module score computed in Scanpy over eight MHC class II genes in tumor cells and averaged per patient: negative or low if the mean patient score was ≤ 0.02 (n = 28), intermediate if > 0.02 and ≤ 0.1 (n = 5), and positive if > 0.1 (n = 9). A low module score is a relative measure and does not indicate absence of MHC class II transcripts. Cell type annotations and cluster assignments were performed as previously described. Spearman rank correlations were calculated between normalized cell type frequencies. All statistical tests were two-tailed.

### Cell lines

The lentiviral packaging cell line Lenti-X, human melanoma cell line SKMel37 and all other SKMel37 derived knockout cell lines were cultured in DMEM supplemented with 10% FBS, 1X Pen-Strep, 1X L-Glutamine and HEPES buffer. All cell lines were incubated at 37°C and 5%CO_2._

SK-MEL-37 and A375 were obtained from ATCC. All cell lines were tested for mycoplasma monthly while in culture.

### Generation of SK-MEL-37 knockout and reporter cell lines

NY-ESO-1 knockout, HLA-DP knockout, and NY-ESO-1/HLA-DP double knockout SK-MEL-37 lines were generated by CRISPR-Cas9 ribonucleoprotein (RNP) nucleofection. Cas9 RNP was prepared as described above for T cells, by combining equal volumes of 80 µmol/L tracrRNA (IDT) and 80 µmol/L crRNA targeting in duplex buffer (IDT), heating to 95 °C for 5 minutes and allowing the mixture to cool slowly, and then mixing the resulting 40 µmol/L duplexed RNA with an equal volume of 24 µmol/L Cas9 protein (IDT) and 1/20 volume of 400 µmol/L Cas9 electroporation enhancer (IDT). SK-MEL-37 cells were nucleofected with RNP on a Lonza 4D nucleofector. Double knockout lines were generated by sequential knockout.

For real-time imaging experiments, target lines were transduced with a lentivirus encoding mCherry (TWIST) prepared as described above for TCR lentivirus. mCherry-positive cells were sorted on and maintained in complete DMEM; mCherry expression was confirmed by flow cytometry immediately before each assay.

### Peptides

PepTivator NY-ESO-1 (Miltenyi biotech; 130-095-380) is a pool of lyophilized peptides, consisting mainly of 15-mer sequences with 11 amino acids overlap, covering the complete sequence of the human NY-ESO-1 protein. This peptide was reconstituted in sterile water as per manufacturer instructions. All other peptides for epitope mapping and minimal epitope detection were synthesized by Elim Biopharma. Lyophilized peptides were reconstituted in DMSO.

### Mice

MITRG (M-CSFʰ/ʰ IL-3/GM-CSFʰ/ʰ TPOʰ/ʰ Rag2⁻/⁻ Il2rg⁻/⁻) and MISTRG (M-CSFʰ/ʰ IL-3/GM-CSFʰ/ʰ hSIRPAᵗᵍ TPOʰ/ʰ Rag2⁻/⁻ Il2rg⁻/⁻) mice were obtained from Regeneron through a materials transfer agreement and maintained as previously described.^38^ F1 mice heterozygous for hSIRPAᵗᵍ were used for experiments. In some of our initial experiments, MI(S)TRG newborn pups were irradiated sublethally (80 rads) within 2 days of birth. But, irradiation was stopped in subsequent experiments because engraftment of human HSCs was observed to be similar with or without irradiation and mice survived longer because they were less anemic when they were not irradiated. Next, they were transplanted with human CD34+ hematopoietic stem and progenitor cells isolated from human fetal liver as described previously^38^. After 7-9 weeks, these mice were bled, and hematopoietic stem cell engraftment was checked by measuring the percentage of human CD45+ cells by flow cytometry. Mice in which human CD45+ cells represented at least 10% of total CD45+ populations (both human and mouse combined) were used for further study.

All animal experimentations were performed in compliance with Institutional Animal Care and Use Committee protocol #202200033.

Male and female mice were used and balanced between experimental groups, and mice were 9-12 weeks of age at the time of tumor implantation. SK-MEL-37 was derived from a male melanoma patient. Mice were assigned to treatment groups on the basis of tumor burden, such that all groups had similar average tumor volumes prior to T cell transfer. Group sizes are given in the figure legends. Mice were euthanized when they developed large tumors or anemia as occurs in MISTRG mice due to loss of murine derived red blood cells by human bone marrow engraftment.

### T cell transduction

On day 0, CD4+ T cells from healthy donors, who provided informed consent on an IRB-approved protocol, were thawed from liquid nitrogen stocks and stimulated with anti-CD3/anti-CD28 microbeads at a 3:1 bead:cell ratio (Dynabeads, Invitrogen) in CTL supplemented with IL2 (50 U/mL), IL7 (5 ng/mL) and IL-15 (5 ng/ml) for 2 days. Also, on day 0, Lenti-X cells (Clontech) were transiently transfected with each TCR plasmid, as well as psPAX2 (Addgene plasmid no. 12260) and pCMV-VSV-G (Addgene plasmid no. 8454) packaging plasmids to prepare the lentivirus. TCR plasmids were prepared by cloning paired TCRα/β chains into a lentiviral expression vector as described previously.^15^

On day +2, activated CD4 T cells were de-beaded using a STEM-Cell magnet and nucleofected with CRISPR-Cas9 ribonucleoprotein (Cas-9 RNP) to knockout the first exon of endogenous TCR alpha and beta chain’s constant region. The Cas-9 RNP was prepared by taking equal volumes of 80 µmol/L TracRNA (IDT) and 80 µmol/L TrbcRNA (IDT) combined with 80 µmol/L of the tracrRNA AGAGTCTCTCAGCTGGTACA in duplex buffer (IDT), mixing well and incubating in a heating block at 95 °C for 5 minutes and allowing the mixture to slowly cool. The resulting 40 µmol/L duplexed RNA was mixed with an equal volume of 24 µmol/L Cas9 protein (IDT) and 1/20 volume of 400 µmol/L Cas9 electroporation enhancer (IDT) and incubated at room temperature for 15 minutes prior to nucleofection.

Cells were nucleofected using a Lonza 4D nucleofector in 100 µL of buffer P3 using program EO-115. Cells were allowed to rest for 4 hours in CTL media prior to lentiviral transduction. Lentiviral supernatant was harvested from Lenti-X cultures, filtered using 0.45-µm polyethersulfone (PES) syringe filters (Millipore), and added to activated T cells in a 24-well tissue culture plate. Polybrene (Millipore) was added to a final concentration of 4.4 µg/mL, and cells were centrifuged at 800 x g and 32 °C for 90 minutes. Sixteen hours later, viral supernatant was replaced with fresh CTL containing IL2 (50 IU/mL), IL7 (5 ng/mL) and IL-15 (5 ng/ml). Half-media changes were then performed every 48 to 72 hours using CTL containing IL2, IL7 and IL-15. Transduced T cells were assessed for transduction efficiency using antibodies specific to CD3 and grown in a rapid expansion protocol described previously^15^ for 10 to 13 days prior to conducting immune assays. To confirm the antigen specificity of the TCR-T cells, CD4 T cells were co-cultured with B-LCL lines pulsed with NY-ESO peptide pools. Specificity was assessed by intracellular cytokine staining for IFN-γ or ELISA for IFN-γ, as indicated.

### B-LCLs-T cells co-culture

B-LCLs (APC) were pulsed with NY-ESO-1 peptide pool at 1 µg/mL for 4 hours at 37 °C, then washed to remove unbound peptide. In parallel, LCLs left unpulsed served as negative controls to establish background reactivity or alloreactivity. NY-ESO-1 TCR-expressing CD4 T cells were co-cultured with peptide-pulsed or control LCLs at a ratio of 2:1 in 96-well round-bottom plates. Golgi Plug was added at the start of coculture to retain intracellular cytokines, and plates were incubated for 5 hours at 37 °C. T cells stimulated with 1X cell stimulation cocktail (eBioscience) and T cells cultured alone served as positive and negative controls, respectively. After coculture, cells were stained with a fixable viability dye (Cat No.) and surface antibodies against CD3 (cat no), CD4 and then fixed and permeabilized and stained intracellularly for IFN-γ. All staining steps were performed at 4 °C in the dark according to the manufacturer’s instructions.

### HLA restriction determination of NY-ESO TCRs

HLA restriction of NY-ESO TCRs was determined using a panel of B-LCL lines expressing defined HLA class II alleles (research cell bank, FHCC) shared with the patient from which the TCR was isolated.^20^ The patient’s HLA class II genotype (HLA-DRB1*01:01/03:01, HLA-DQB1*02:01/05:01, HLA-DPB1*04:01/04:02) was determined by running xHLA^43^ on whole exome sequencing.^20^ TCR-transduced CD4 T cells were co-cultured with either NY-ESO peptide pulsed B-LCL lines or autologous B cells (positive control) and IFN-γ production was measured by intracellular cytokine staining (ICS).

### Epitope mapping of the TCR

For epitope mapping, a library of 21 overlapping 20-mer peptides (12-amino acid overlap) spanning the full-length NY-ESO-1 protein was synthesized. Individual peptides were tested for recognition by the HLA-DP restricted NY-ESO specific TCR. CD4 T cells and peptide (1 ug/ml) pulsed HLADP0401^+^ B-LCL line were plated at indicated ratio in 96-well round-bottom plates in the presence of brefeldin A and golgi plug for 5 hours. 1x cell stimulation cocktail was kept as the positive control, and unrelated TCR-transduced T cells from the same donor as a control effector population. Recognition was measured as IFN-γ production by CD4 T cells by ICS. A minimal epitope corresponding to NY-ESO-1 residues 161–172 (WITQCFLPVFLA) was identified as required for recognition by this TCR; the same sequence is present in the related CTAG1A, CTAG1B and CTAG2 (LAGE-1a) genes.

### Alanine Scanning for determining minimal epitope

A panel of single-point alanine substitution variants was designed based on the wild-type (WITQCFLPVFLA) epitope. For functional evaluation, irradiated DP0401^+^ PBMCs as antigen-presenting cells (APCs) were pulsed with either WT or alanine-substituted peptides for 4 hours at 37 °C. APCs were then co-cultured with NYESO TCR-transduced CD4 T cells at an APC-to-T cell ratio of 1:4 in a 96-well round-bottom plate in the presence of Golgi plug. After 5 hours, T cell activation was quantified by measuring the percentage of IFN-γ expressing CD4 T cells by flow cytometry. Recognition was assessed at a high and low peptide concentration, and contact residues were defined as positions where recognition at the high concentration was less than 60% and at the low concentration less than 20% of the wild-type peptide response. Proteome scanning was performed using the ScanProsite tool.^44^ The resulting recognition motif, W-x-x-Q-C-F-L-P-x-x-x, with critical residues at positions 4–8, was returned only by NY-ESO-1 (CTAG1B) and the related CTAG1A and CTAG2 (LAGE-1a) cancer-testis antigens.

### Direct antigen recognition assays

For direct recognition assays, TCR-transduced CD4 T cells were cocultured with SK-MEL-37 melanoma cells (wild-type, NY-ESO-1 knockout, HLA-DP knockout, or double knockout generated by CRISPR/Cas9) at target to effector ratio of 1:2 and IFN-γ production was measured by ICS.

### Monocyte-derived dendritic cell (moDC) preparation

Autologous leukapheresis samples were washed with serum free RPMI, set to 5–10 × 10⁶ cells/mL and then incubated at approximately 1 × 10⁶ cells/cm² in an appropriately sized tissue culture flask (T150) and incubated for 90 minutes at 37 °C, 5% CO₂. After 90 min, non-adherent cells were discarded, and the flasks were vigorously washed with serum free RPMI and then incubated in serum free RPMI for another 60 mins. The flasks were then vigorously washed again with serum free RPMI, and then the adherent cells were incubated with DC media (freshly prepared). This day is considered Day 0. DC cultures were supplemented with fresh DC media every 3 days. DCs were used in co-culture experiments on day 7 after initial stimulation. In all experiments, flow cytometry was used to phenotype the cells for expression of CD11c, CD14, CD80, CD86, and HLA-DR (all from BD Bioscience) to ensure that the cells were predominantly immature DCs (CD11c+, CD14-, CD80low, CD86+, and HLA-DR+).

### Tumor cell lysate preparation

Whole-cell lysates were prepared from NY-ESO-1–positive, NY-ESO-1 knockout, and HLA-DP knockout SK-MEL-37 melanoma cell lines. Typically ∼5 × 10⁶ cells were harvested, washed, and resuspended in 500–600 µL of 1× PBS to maximize protein concentration. Cells were lysed by six freeze–thaw cycles, consisting of rapid freezing in liquid nitrogen followed by thawing in a 37 °C water bath; no protease inhibitors were included. Total protein concentration of each lysate was determined using a BCA protein assay kit (Thermo Fisher Scientific) according to the manufacturer’s instructions and expressed in µg/mL. Lysates were stored at −80 °C if not used immediately.

### Generation and antigen loading of monocyte-derived dendritic cells

Immature monocyte-derived dendritic cells (moDCs) were generated from an HLA-DP0401+ donor over a 6-day differentiation period. On day 7, immature moDCs were seeded into flat-bottom TC treated 96-well plates at 5-8 × 10³ cells per well in 250 µL of DC medium and loaded with tumor cell lysate at a final concentration of 50-80 µg/mL. Loading was performed in the presence of recombinant human GM-CSF (800 U/mL) and recombinant human IL-4 (800 U/mL), and cells were matured overnight. Surplus moDCs were cryopreserved.

### moDC–T cell co-culture representing indirect antigen presentation

On the day before the co-culture, rapidly expanded NY-ESO CD4 T cells were thawed and rested overnight in CTL medium supplemented with recombinant human IL-2 (15 U/mL). Lysate-loaded, matured moDCs (5 × 10³) were co-cultured with NY-ESO-1 specific CD4⁺ T cells (2.5 × 10⁴). Co-cultures were established directly in the 96-well plates in which the moDCs had been seeded; at setup, half of the DC medium was removed and replaced with an equal volume of CTL medium without added cytokines. The co-cultures were incubated overnight and supernatants were collected for quantification of secreted IFN-γ by ELISA (Thermo fisher).

### Incucyte Killing Assay

To enable real-time tracking of target tumor cells, they were lentivirally transduced with mCherry fluorescent reporter protein. mCherry positive tumor cells were sorted and maintained in complete DMEM. mCherry labeled target tumor cells were seeded at a density of 5 × 10³ cells per well into a flat-bottomed 96-well plate and allowed to adhere overnight. Next day, effector CD4 T cells (NYESO/Control) and freshly prepared monocytes/moDCs (HLADP04 matched with the TCR) were added at T:APC:target ratios of 4:1:1 in a final volume of 200 µL/well of CTL. To dissect the killing pathways involved in T cell mediated killing, neutralizing antibodies were added to the co-culture at assay initiation and maintained throughout imaging. Anti-IFN-γ, anti-TNF-α and anti-FasL were added either individually or in combination at a concentration of 10 µg/mL. Target-only wells served as the baseline control. Plates were placed in the Incucyte S3 live-cell imaging system (Sartorius) at 37 °C and 5% CO₂ and whole wells were imaged every 4 hours over 5 days using the 4X objective.

For quantifying killing, images were analyzed using Incucyte software. Live target cells were quantified as the total Red Object Integrated Intensity (RCU × µm²/well), which integrates red-object fluorescence intensity over object area summed across the well, normalized to the value at hour 0. Each condition was run in triplicates.

### Tumor implantation and adoptive T cell transfer in humanized mice

SK-MEL-37 (HLA-DP-positive) and SK-MEL-37 HLA-DP knockout cells were implanted subcutaneously in opposite flanks. Tumor-bearing mice received adoptive transfer of NY-ESO-1-specific or irrelevant TCR-transduced CD4 T cells at 10^7^ cells 14 days after tumor implantation. Tumor volume was measured by caliper every as indicated. For phenotypic analysis, epitope-tagged T cells were transferred, and tumors were harvested 8 days post-transfer for flow cytometric analysis. 1 × 10⁶ SK-MEL-37 cells in 100µL of 1:1 mixture of PBS and Matrigel were implanted subcutaneously into each flank. Tumor volume was calculated as (length × width²)/2 and measured twice weekly beginning 7-10 days after implantation.

### Processing of tumors

Tumors were excised, minced, and dissociated to a single-cell suspension using the human Tumor Dissociation Kit on a gentleMACS Octo Dissociator with heaters (Miltenyi Biotec) according to the manufacturer’s instructions.

### Single-cell capture and cDNA library preparation for single cell RNA sequencing

Single-cell libraries were generated on the BD Rhapsody Express system (BD Biosciences) using the targeted mRNA and AbSeq Reagent Kit-4 pack, following the manufacturer’s instructions throughout. Single cell suspensions of MISTRG tumors were prepared by enzymatic digestion using human tumor dissociation kit (Miltenyi Biotech) as per manufacturer’s instructions. Each sample was labeled with a unique sample tag from the BD Single-Cell Multiplexing, allowing samples to be pooled and later demultiplexed in silico. Tagged cells were washed in BD Stain Buffer, blocked with Fc receptor blocking reagent, and then labeled with the BD AbSeq Ab-Oligo master mix (3 µL per Ab-Oligo). To enrich for the lymphoid and myeloid populations of interest, the sample was co-stained for 45 min at 4 °C with fluorochrome-conjugated antibodies against human CD3 (clone OKT3), CD4 (RPA-T4), CD8 (RPA-T8), CD19 (HIB19) and CD45 (2D1) (BD Biosciences or BioLegend) and sorted by flow cytometry.

Sorted populations were counted and pooled to give 20,000 cells in 620 µL of BD Sample Buffer. This suspension was loaded onto a primed BD Rhapsody cartridge and allowed to settle at room temperature, after which washed Cell Capture Beads were loaded and incubated. The cartridge was washed twice with BD Sample Buffer, cells were lysed, and the beads were retrieved, washed, and taken immediately into reverse transcription.

From this point we followed the BD Rhapsody VDJ CDR3, Sample Tag and AbSeq library protocol, which allowed TCR and BCR libraries to be generated alongside the mRNA, AbSeq and Sample Tag libraries from the same beads. A poly-T template-switching oligo (TSO) was included in the reverse transcription reaction so that VDJ recombination events in T and B cells could be captured. Reverse transcription was followed by denaturation, hybridization, Klenow extension (New England Biolabs, #M0212L) and Exonuclease-I treatment.

cDNA libraries were built by two-step nested PCR followed by index PCR, using the BD Rhapsody Targeted mRNA and AbSeq Amplification Kit (#633774) with the BD Rhapsody Immune Response Panel Hs (#633750).

After the first PCR, the longer mRNA, TCR and BCR products were separated from the shorter AbSeq and Sample Tag products by double-sided size selection with Agencourt AMPure XP beads (Beckman Coulter, #A63880). Library concentrations were determined with the Qubit dsDNA HS Assay Kit (Thermo Fisher Scientific, #Q32851), and second-PCR products were diluted according to the protocol’s recommendations before index PCR. Final libraries were assessed for quality and quantified using Agilent 2200 TapeStation with High Sensitivity D5000 ScreenTape (Agilent). Sequencing was performed on a Novaseq SP flow cell at 75 × 225 and after demultiplexing, sample tags, mRNA, and antibody sequencing reads were trimmed to 75nt and TCR and BCR VDJ libraries were kept at 225 nt and FASTQ files were uploaded to the BD Rhapsody analysis pipeline () with refined cell calling disabled.

### Single-cell RNA sequencing analysis

Gene and surface protein expression matrices were analyzed using the Scanpy version 1.11.3^45^ toolkit in Python. Low quality cells were removed (designated as cells with fewer than 10 expressed genes or 100 total counts). Doublets were removed using the Scrublet package.^46^ Gene expression counts were log-normalized, and protein counts were transformed using centered log-ratio (CLR) normalization.

Principal component analysis was performed on the gene expression data, and a UMAP projection was generated based on the top 20 principal components. High-level unsupervised Leiden clustering was performed (using a resolution parameter of 0.7), and the broad clusters were manually labeled based on expression of classical myeloid and T cell marker genes. Myeloid cells and T cells were then separately subclustered, using Leiden clustering with a resolution parameter of 1. Differentially expressed genes for subclusters were identified with rank_genes_groups in Scanpy under its default settings, which apply a t test with Benjamini-Hochberg adjustment; these marker tables were used for cluster annotation only.

### Estimation of HLA-DP04 haplotype prevalence

Frequencies of the DPA1*01:03∼DPB1*04:01 and DPA1*01:03∼DPB1*04:02 haplotypes were taken from the NMDP/Be The Match registry haplotype frequency dataset,^34^ which reports two-locus DPA1∼DPB1 haplotype frequencies for registry volunteers grouped into 26 population categories: 5 broad categories and 21 more finely resolved categories nested within them.

Because the TCR is restricted by the DP0401/DP0402 heterodimer, both the DPA1*01:03 alpha chain and a DPB1*04:01 or *04:02 beta chain are required; for each population category the frequencies of the two qualifying haplotypes were therefore summed to give the frequency f of any TCR-compatible haplotype, and the proportion of individuals carrying at least one copy was calculated as 1 − (1 − f)² under Hardy–Weinberg equilibrium.

### Statistical analysis

Comparisons between two groups used the two-tailed Mann–Whitney U test or Welch’s unequal-variance t test, as specified in each figure legend. Spearman rank correlation was used to assess associations between cell-type frequencies, with permutation P values, and overall survival was compared by log-rank test. All tests were two-tailed and P < 0.05 was considered statistically significant. Significance is shown as asterisks on most panels, with exact P values reported in the figure source data; the test used and any correction applied are stated in the corresponding figure legend. Mann–Whitney U tests for the Figure 5 panels used the exact null distribution. Corrections for multiple comparisons were not applied to the P values annotated on the figure panels. Benjamini-Hochberg adjusted values are reported where a family of tests is interpreted together: the specificity screen in Fig. S2D and the 220-gene comparison in Fig. 6J. Where a panel derives from a single experiment with technical replicates, the legend states the number of wells and notes that the resulting P values describe within-experiment variation. For scRNAseq analysis of myeloid cells expression, each gene, the probability of detection per cell was modeled as a function of that cell’s sequencing depth by logistic regression across all myeloid cells; each mouse is summarized as log2 of its observed detection fraction divided by the fraction expected at its own cells’ depths, so that 0 denotes exactly the depth-predicted rate. Module score is the mean of the five per-gene values. Points are per-mouse values, horizontal bars are group means, and marker area is proportional to the number of myeloid cells contributing to that mouse. All comparisons aggregate to the mouse before testing. Myeloid comparisons (panels h–j) included only mice with at least 5 myeloid cells (NY-ESO n = 3; control n = 5). The CD8 module comparison in Fig. 6F used n = 6 NY-ESO and n = 5 control mice, because one control mouse yielded fewer than two CD8 T cells. The per-mouse comparison of WPRE+ cells in Fig. 6E used n = 6 mice per group. The two modules plotted in Fig. 6E were specified in advance of the comparison, and each was scored by standardizing every constituent gene across the twelve per-mouse values and averaging the standardized values over the genes of that module present on the targeted panel (7 of 7 proliferation genes and 10 of 10 co-inhibitory genes); the two module scores were compared between arms by two-tailed exact Mann-Whitney U. Module z-scores in Fig. 6D were computed per gene by subtracting the mean across all cells and dividing by the standard deviation among cells with nonzero expression, then averaging over the genes of each signature present on the targeted panel. Analyses were performed in GraphPad Prism (GraphPad Software), R and Python.

### Study approval

All animal experiments were performed with the approval of the Fred Hutchinson Cancer Center Institutional Animal Care and Use Committee. Peripheral blood and leukapheresis products from de-identified healthy donors were obtained with written informed consent from Bloodworks northwest and StemCell. Tumor tissue and blood from the index melanoma patient were obtained with written informed consent under protocol FH2643, approved by the Fred Hutchinson Cancer Center Institutional Review Board. Human CD34+ hematopoietic stem and progenitor cells were isolated from human fetal liver obtained from deidentified donors that signed informed consent. All studies were conducted in accordance with the Declaration of Helsinki and the Belmont Report.

## Data Availability

Raw and processed targeted single-cell RNA sequencing data generated in this study, together with the paired AbSeq, sample-tag and TCR data, the cell-to-sample metadata, and the custom targeted-panel reference including the WPRE amplicon, are deposited in the Gene Expression Omnibus under accession GSE348872, and will be made available on publication of peer reviewed paper or on reasonable request. The melanoma cohort single-cell RNA sequencing data analyzed here were previously published and are available in GEO at GSE198265. The breast cancer single-cell RNA sequencing data from Bassez et al.^32^ are available at EGA under EGAD00001006608 through controlled access requiring data access committee approval. Any additional information required to reanalyze the data reported in this paper is available from the lead contact upon request.

## Code Availability

All original code is deposited at Zenodo 10.5281/zenodo.23045730, and will be made available on publication of peer reviewed paper or on reasonable request

## Resource Availability

Plasmids use in this work will be deposited at addgene by time of publication. Modified cell lines will be available on request. MISTRG mice require a materials transfer agreement with Regeneron.

Lead contact: Joshua Veatch

## Generative AI use statement

Claude Opus 4.6–5.0 was used to assist with data analysis for Figs. 1, S1, 5, and 6, and with data visualization and figure organization throughout. The authors (C.S.) reviewed and verified all code developed with AI assistance and the resulting analyses and confirmed reproducibility. Claude Opus 4.6–5.0 and ChatGPT 5.5–6 were also used to assist with literature review, manuscript organization, language editing, and feedback on manuscript drafts. The authors developed the study’s concepts, directed the analyses, interpreted the findings, and critically revised and approved the final manuscript. The authors take full responsibility for the accuracy and integrity of the work.

## Acknowledgements

We would like to thank Regeneron for providing MISTRG mice for this study, and Stanley Riddell and Shivani Srivastava for feedback on this manuscript, and to patients who provided research samples and their families. This work was supported by grants from the NCI (R37CA292663), Melanoma research foundation (1143984), Melanoma Research alliance (1263967), and Washington Research Foundation to J.R.V. as well as the Fred Hutchinson Cancer Center consortium grant P30 CA015704.

## Author Contributions

G.R.: Conceptualization, Methodology, Validation, Investigation, Visualization, Writing – original draft, Writing – review & editing. M.R.: Conceptualization, Methodology, Investigation, Writing – review & editing. J.B.: Conceptualization, Methodology, Investigation, Writing – review & editing. C.S.: Software, Formal analysis, Validation, Visualization. S.L.: Resources. A.M.: Methodology, Investigation. C.W.: Investigation. K.H.: Investigation. M.W.: Investigation. J.S.: Investigation. N.S.: Investigation. M.I.: Investigation. K.M.M.: Investigation. E.N.: Conceptualization, Writing – review & editing. A.R.: Conceptualization, Methodology, Resources. J.R.V.: Conceptualization, Methodology, Validation, Visualization, Supervision, Funding acquisition, Writing – original draft, Writing – review & editing.

---

**Figure S1.**
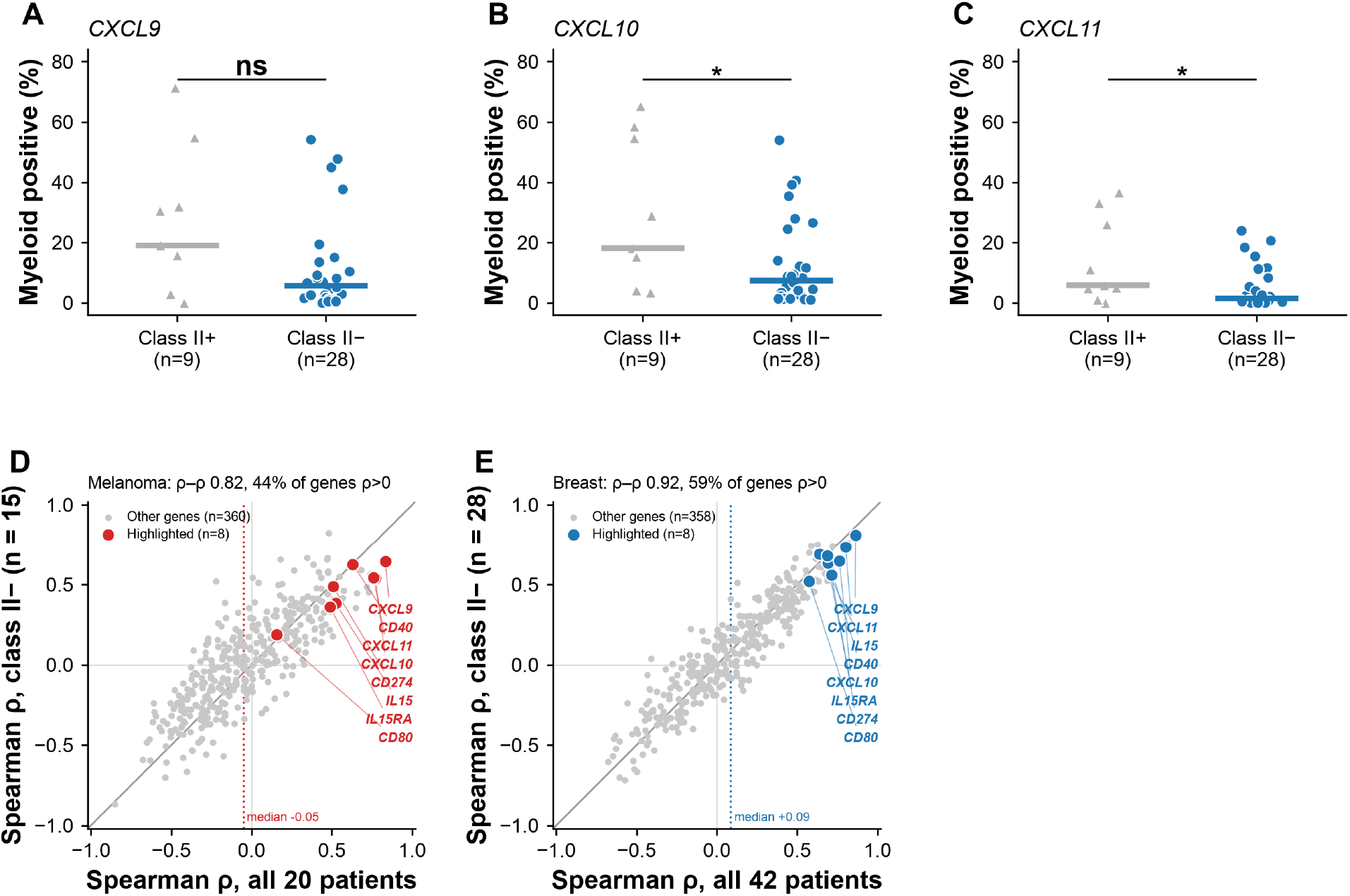
Myeloid chemokine detection by tumor MHC class II status, and gene-level comparison of full-cohort and class II-negative correlations. (A–C) Myeloid detection of (A) CXCL9, (B) CXCL10 and (C) CXCL11 in the breast cancer cohort, compared between MHC class II-positive (n = 9) and class II-negative (n = 28) tumors; the 5 intermediate cases are excluded. Points are patients, horizontal bars are medians, and values are raw percent of myeloid cells positive. (D, E) Spearman ρ between CXCL13+ CD4 T cell frequency and myeloid detection of each gene on the panel, computed in all patients (x axis) against the class II-negative subset alone (y axis), for (D) the melanoma cohort (368 genes; 20 versus 15 patients) and (E) the breast cohort (366 genes; 42 versus 28 patients). Genes carried through the main figures are labeled; the diagonal is the line of equality and the dotted line marks the median ρ across all genes. Genes detected in fewer than 5 cells, or in every cell, are omitted because a depth model cannot be fitted. Myeloid gene detection is expressed as log2 of the observed detection rate over the rate expected from each cell’s sequencing depth. Group comparisons are by Mann-Whitney U test; all tests are two-tailed. Significance is shown on panels (A–C) as asterisks (*P < 0.05, **P < 0.01, ***P < 0.001, ****P < 0.0001; ns, not significant). Panels (D) and (E) compare Spearman ρ values between cohorts. No correction for multiple comparisons was applied.

**Figure S2.**
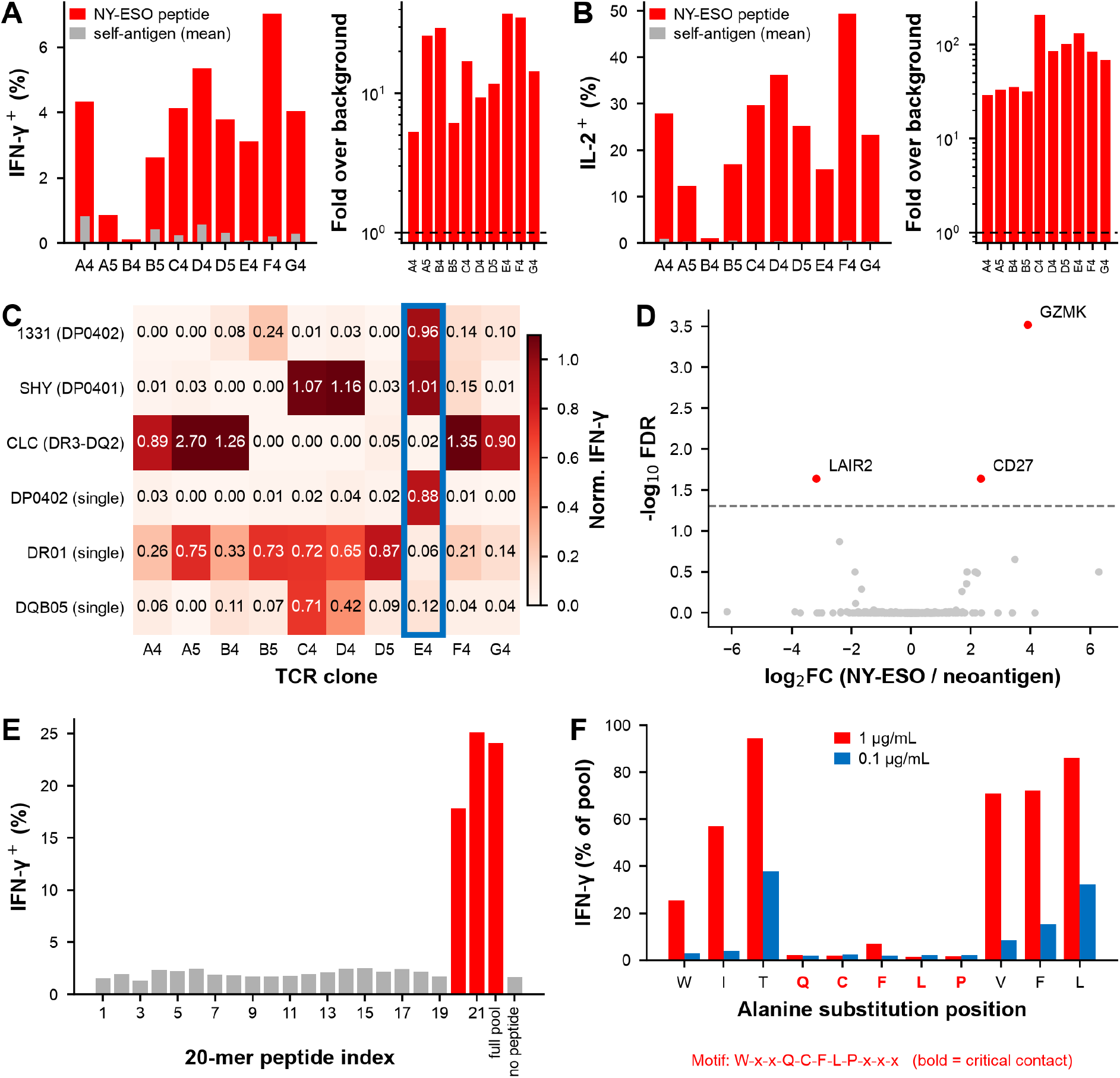
Characterization of NY-ESO-1-specific CD4 TCRs: antigen screen, HLA restriction, differential expression, and epitope mapping. (A, B) Reactivity of candidate shared-antigen TCRs against individual antigen peptide pools, assessed by intracellular IFN-γ (A) and IL-2 (B) staining following coculture with autologous B cells; all recovered TCRs recognized NY-ESO, and for TCRs with multiple candidate α-chain pairings, specificity was recovered for one of the two alternate pairings tested. (C) HLA restriction of each NY-ESO-specific TCR determined using B-lymphoblastoid cell lines and single-allele lines; the lead TCR (E4) is restricted to HLA-DPB1*04:01/04:02. (D) Differential gene expression between NY-ESO-1-specific and neoantigen-specific CD4 T cells in the primary tumor, highlighting GZMK and CD27. (E) Epitope mapping using 21 overlapping 20-mer peptides spanning NY-ESO, with the full peptide pool and a no-peptide control shown as the last two bars; two adjacent C-terminal peptides drive recognition, localizing a minimal epitope corresponding to NY-ESO-1 residues 161–172 (WITQCFLPVFLA), whose contact residues were subsequently defined by alanine scanning (Fig. S2F). (F) Alanine-scanning mutagenesis of the NY-ESO epitope; the recognition motif W-x-x-Q-C-F-L-P-x-x-x is shown, with glutamine, cysteine, phenylalanine, leucine, and proline (positions 4–8) as critical residues. A proteome-wide search for this motif returned only NY-ESO (CTAG1B) and the related CTAG1A and CTAG2 (LAGE-1a) cancer-testis antigens.

**Figure S3.**
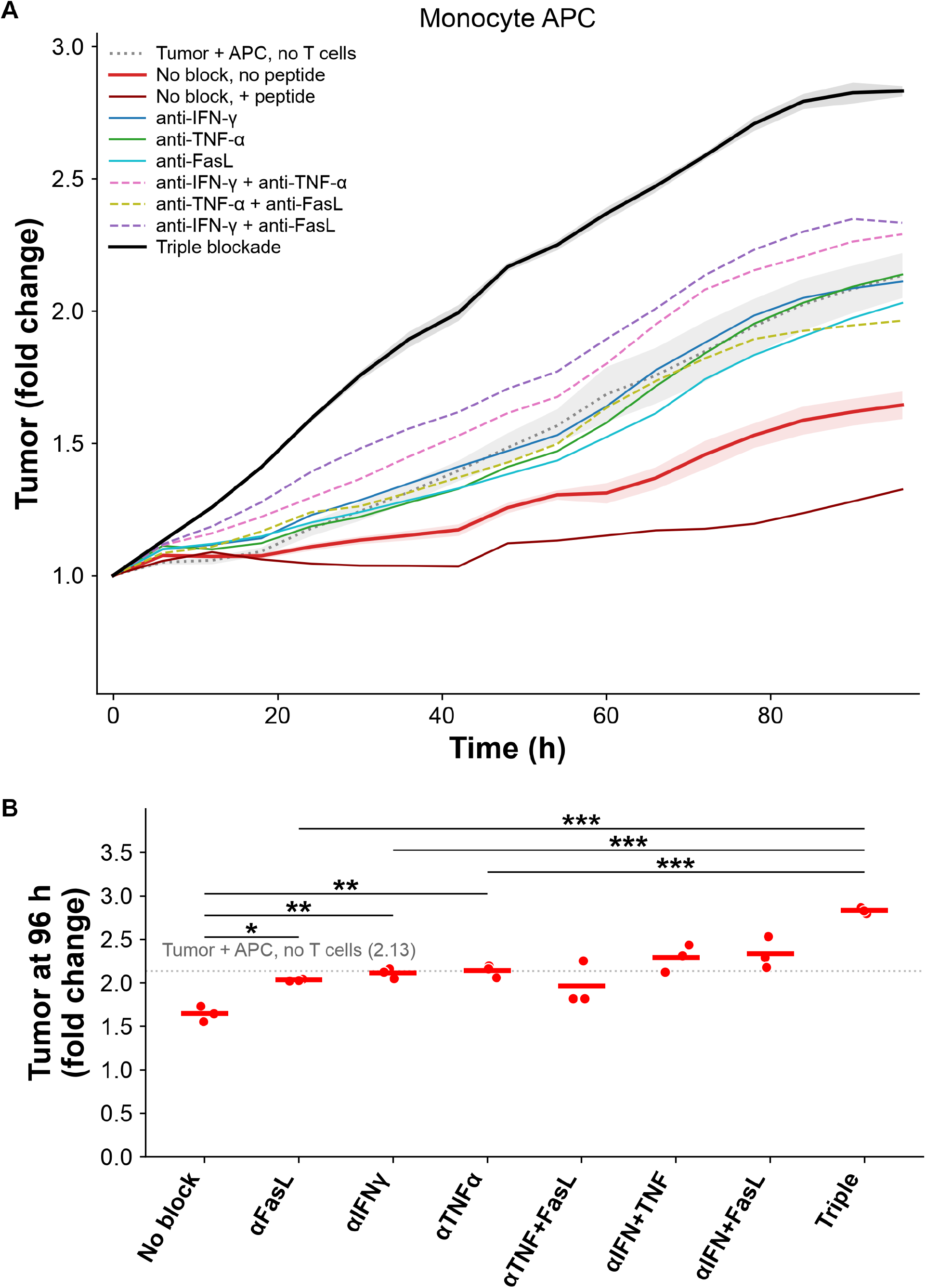
Indirect control of tumor growth with peripheral blood monocytes as APCs. (A) Time course of the monocyte indirect control assay under blockade of IFN-γ, TNF-α and FasL, alone and in combination, with no-block and no-T-cell (tumor + APC) references, over 96 h. (B) Endpoint (96 h) tumor fold change across the same single and combined blockade conditions; dotted line = tumor + APC, no T cells reference (2.13). Horizontal bars are arithmetic means; n = 3 replicate wells per condition from a single experiment. In (B), endpoint tumor fold change was compared between conditions by two-tailed Welch’s t test on n = 3 replicate wells per condition without correction for multiple comparisons (*P < 0.05, **P < 0.01, ***P < 0.001, ****P < 0.0001; ns, not significant).

**Figure S4.**
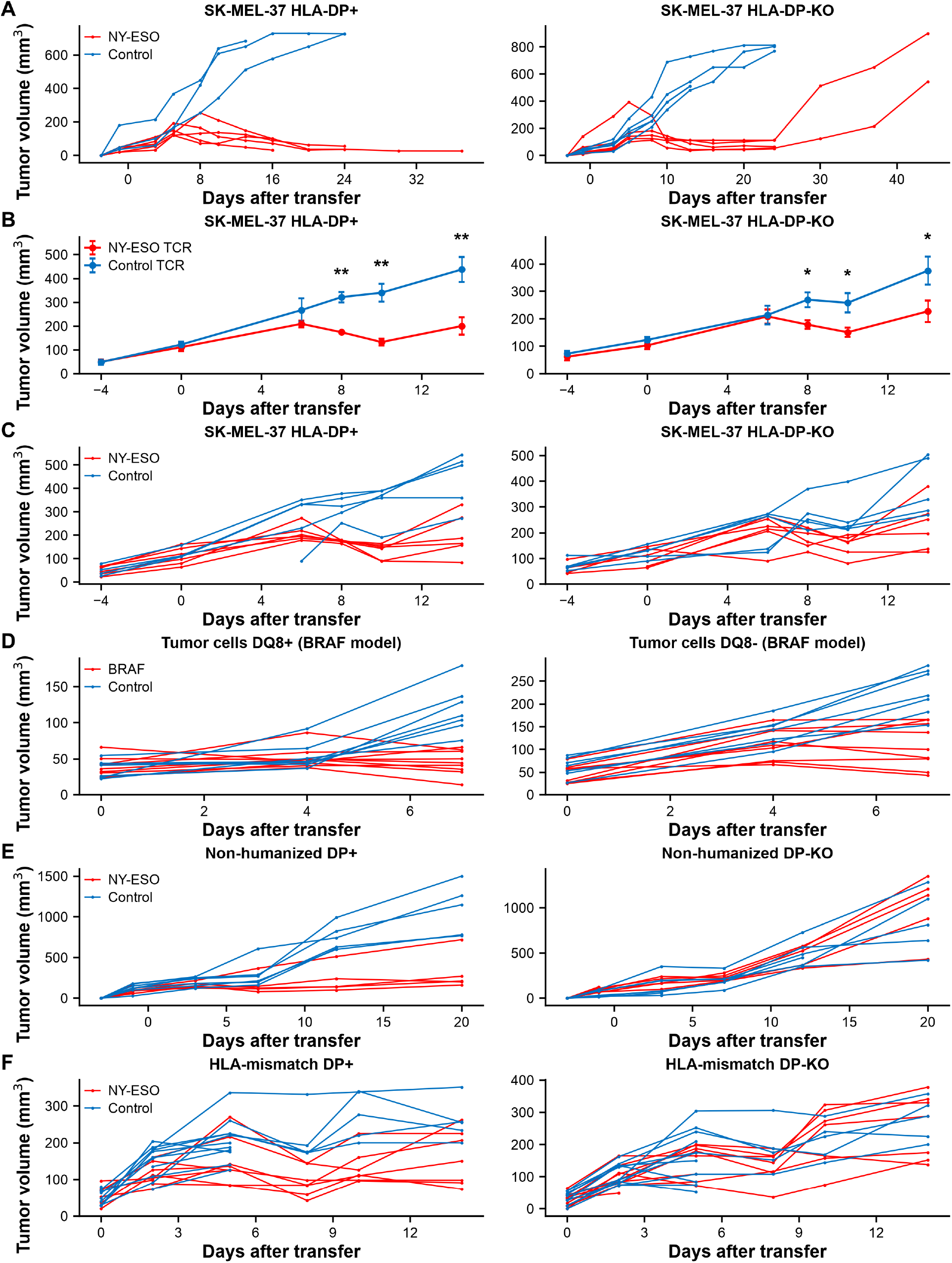
Individual-mouse tumor curves, later timepoints, and an independent donor repeat for the arms in Fig. 4. In every row the left panel is the HLA-DP-expressing (or HLA-DQ8-positive) flank and the right panel the HLA-DP knockout (or HLA-DQ8-negative) flank of the same animal; control TCR in blue, NY-ESO-specific or BRAF-V600E-specific TCR in red. (A) Individual tumor trajectories for the humanized HLA-DP04 experiment in Fig. 4C, over the full observation period. (B, C) An independent repeat of that experiment in a separate cohort using different HSC and CD4 T cell donors, shown as (B) mean ± SEM per timepoint and (C) individual tumor trajectories (n = 6 NY-ESO TCR, 5 control TCR; In (B), asterisks denote nominal significance for the difference in tumor volume between arms at that timepoint by Welch’s t test, applied independently at each timepoint with no correction for multiple comparisons (*P < 0.05, **P < 0.01, ***P < 0.001, ****P < 0.0001; ns, not significant). (D) Individual trajectories for the HLA-DQ8 BRAF-mutant experiment in Fig. 4D. (E) Non-humanized MISTRG (Fig. 4E). (F) HLA-DP04-mismatched HSC (Fig. 4F). Mice censored with small tumors died due to anemia from engraftment of human HSC precluding murine red blood cell formation.

**Figure S5.**
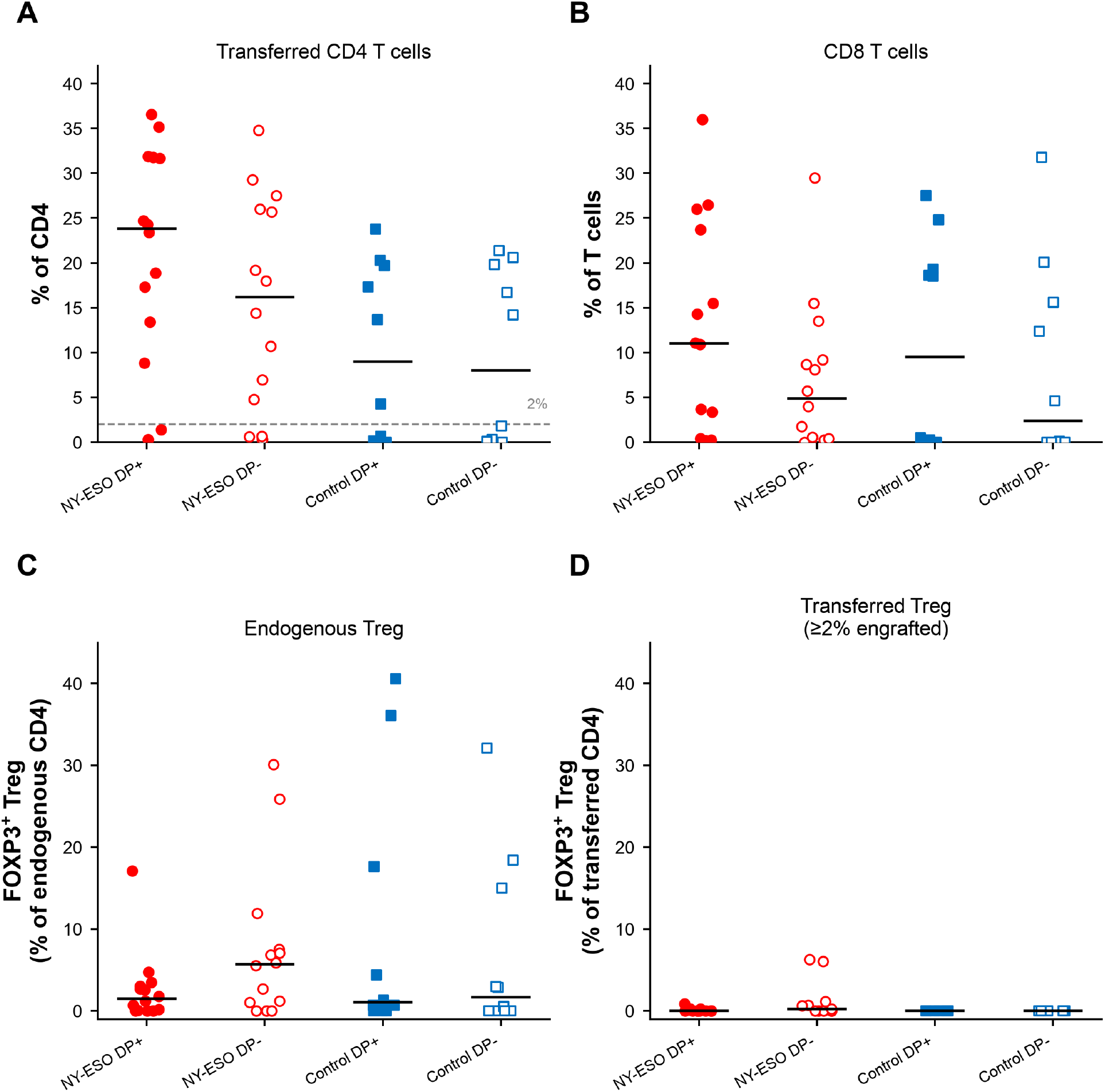
Engraftment of human immune cells in SK-MEL-37 tumor xenografts 8 days following T cell transfer. A) NGFR+ transferred T cells as a fraction of total CD4+ T cells (NGFR+ T cells were 50% of infusion product) B. CD8 T cells as a fraction of CD3+ T cells. C. FoxP3+ cells as a fraction of CD3+ CD4+ NGFR-CD4 T cells. D. FoxP3+ cells as a fraction of NGFR+ CD4+ T cells, in tumors with ≥2% engraftment of transferred cells. Points are individual tumors, horizontal bars are medians.

**Figure S6.**
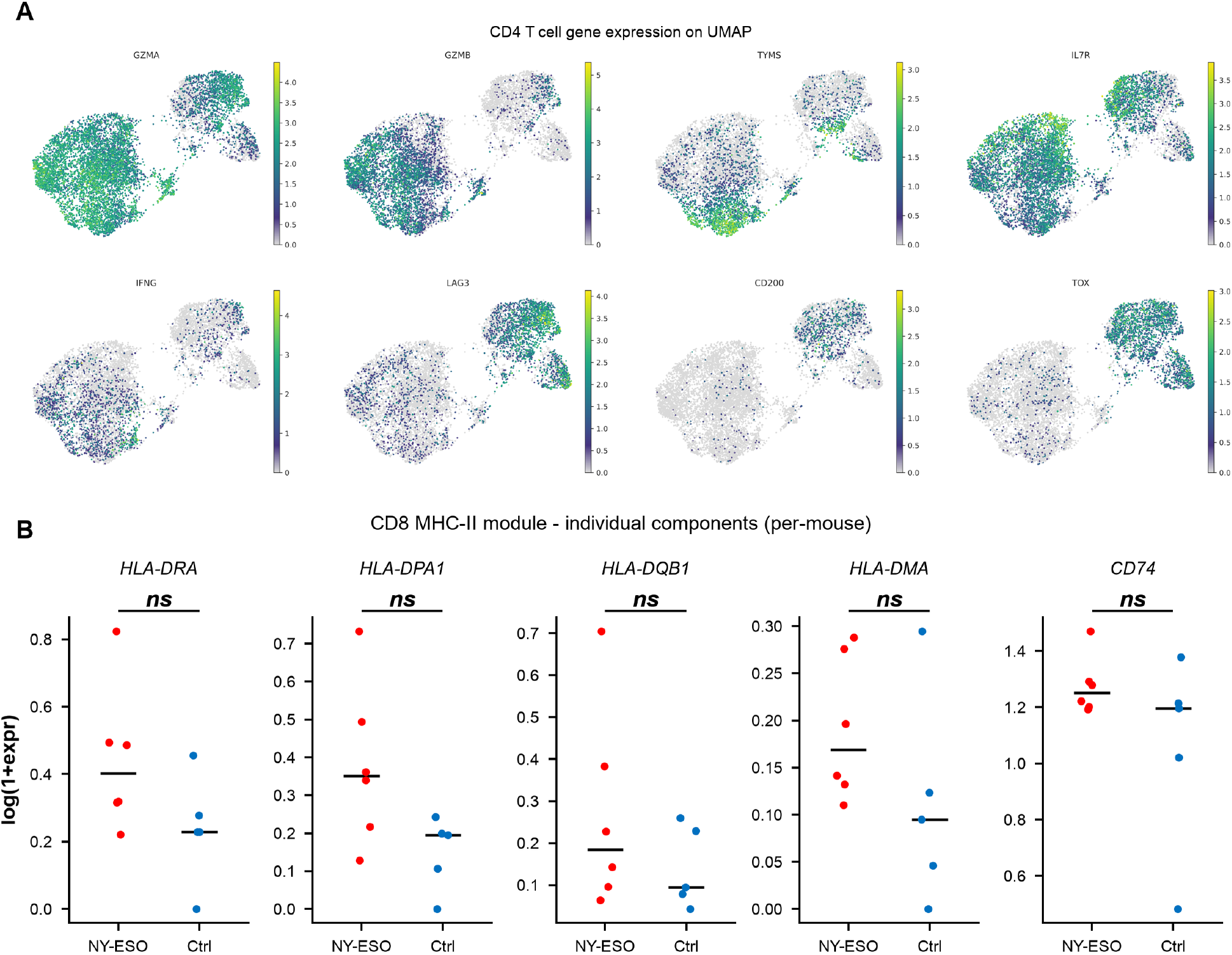
Human CD4 T cell gene expression on the MISTRG single-cell UMAP, and individual components of the CD8 MHC class II module. (A) Expression of GZMA, GZMB, TYMS, IL7R, IFNG, LAG3, CD200 and TOX projected onto the CD4 T cell UMAP embedding shown in Fig. 6C; color indicates normalized expression and is scaled independently for each gene. (B) Per-mouse mean expression of the individual CD8 MHC class II module genes (HLA-DRA, HLA-DPA1, HLA-DQB1, HLA-DMA and CD74) in NY-ESO versus control mice, as log(1+expression); horizontal bars are group medians. Statistics: in (B), per-mouse mean expression was compared between NY-ESO and control mice by two-tailed Mann-Whitney U, uncorrected for multiple comparisons, with significance shown on each panel (*P < 0.05, **P < 0.01, ***P < 0.001, ****P < 0.0001; ns, not significant).

**Table S1.** Melanoma patient cohort characteristics and tumor MHC class II status (Related to Fig. 1).

| Patient ID | Sex | Age | Site of harvest | Treated with TIL ? | Best response to TIL | Prior therapies | Subsequent therapies | BRAF status | Survival (years) | Deceased? | Tumor MHC class II by IHC (pathologist read) | MHC class II group |
| --- | --- | --- | --- | --- | --- | --- | --- | --- | --- | --- | --- | --- |
| x176 | M | 54 | ABD mass | No | — | nivolumab | surgery | BRAF V600E | 6 | No | Heterogenous, weak expression within tumor, melanoma cells (big nuclei, prominent nucleoli, lots of cytoplasm) are often negative but there are admixed lymphocytes that stain positive | Negative |
| x197 | M | 52 | L iliac node/L thigh | Yes | CR | ipilimumab | TIL with CR | BRAF V600E | 6 | No | Positive | Positive |
| x198 | M | 63 | Right thigh | Yes | CR | ipilimumab | TIL with CR | wildtype | 6 | No | Positive | Positive |
| x205 | F | 57 | Right breast | No | — | mage a3 vaccine, ipilimumab, intralesional il-2 | surgery, pembrolizumab (? Response) | wildtype | 6 | No | Negative | Negative |
| x215 | F | 30 | Rt back | Yes | PD | interferon, ipilimumab+IL-21, dacarbazine, pembrolizumab, gamma knife, dabrafenib/trametinib | TIL, dabrafenib/trametinib, ipilimumab/nivolumab, surgery | Mutant | 5 | No | Negative | Negative |
| x222 | M | 59 | Back | No | — | none | — | wildtype | 1.5 | Yes | Negative | Negative |
| x226 | F | 40 | Scalp | No | — | Ipilimumab | dabrafenib/trametinib, nivolumab | Mutant | 1.24 | Yes | Heterogenous, weak expression within tumor, melanoma cells (big nuclei, prominent nucleoli, lots of cytoplasm) are mostly negative but there are admixed melanophages and other inflammatory cells that stain positive | Negative |
| x229 | F | 32 | Rt inguinal node | No | — | Ipilimumab+intralesional IL-2, pembrolizumab | — | mutant | 2.44 | Yes | Heterogenous, mostly positive staining within the tumor. | Positive |
| x231 | M | 55 | Gallbladder | Yes | PD | GVAX, ipilimumab | pembrolizumab | wildtype | 2.1 | Yes | Negative | Negative |
| x257 | M | 53 | Mesentery | No | — | dacarbazine, ipilimumab, PD-1 | vaccine + nivolumab | wildtype | 6 | No | Negative | Negative |
| x266 | M | 58 | Lf chest wall | No | — |  | ipilimumab + nivolumab | wildtype | 0.42 | Yes | Negative | Negative |
| x275 | M | 46 | R retroperitoneum | No | — | ipilimumab, radiation | nivolumab, dabrafenib/trametinib | BRAF V600E | 0.3 | Yes | Negative | Negative |
| x330 | F | 62 | Left lower flank/upper buttock subcutaneous tissue | No | — | Ipilimumab, then pembrolizumab with progression on both | had delayed pembro response | wildtype (NRAS mutant) | 5.5 | No | Negative | Negative |
| x353 | M | 61 | Lt axillary node | No | — | High dose IL-2, ipilimumab, pembrolizumab, dabrafenib/trametinib, radiation | — | BRAF V600K | 0.33 | Yes | Negative | Negative |
| x368 | M | 40 | Right Axilla | No | — | none | adjuvant ipilimumab, then ipi/nivo after progression | braf v600E | 4 | No | Heterogenous, weak expression within tumor, melanoma cells (big nuclei, prominent nucleoli, lots of cytoplasm) are mostly negative but there are admixed melanophages and other inflammatory cells that stain positive | Negative |
| x372 | M | 31 | right medial thigh | Yes | PD | interferon | ipi+radiation | wildtype (NRAS mutated) | 2.71 | Yes | Heterogenous, mostly positive staining within the tumor. | Positive |
| x374 | F | 52 | Brain | No | — | ipi/nivo | ipi+radiation, imatinib | wildtype (Kit mutated) | 1.7 | Yes | Negative | Negative |
| x392 | M | 54 | Rt Axilla | No | — | dabrafenib/trametinib, nivolumab | — | Mutant | 0.8 | Yes | Positive | Positive |
| x422 | F | 59 | r thigh | No | — | none | none | wildtype | 1.4 | Yes | Negative | Negative |
| x431 | M | 62 | liver | No | — | tumor lysate vaccine | surgeries | wildtype | 3.2 | Yes | Heterogenous, weak expression within tumor, melanoma cells (big nuclei, prominent nucleoli, lots of cytoplasm) are often negative but there are admixed inflammatory cells that seem to be positive (many plasma cells that are possibly taking up the stain) | Negative |

**Table S2.**
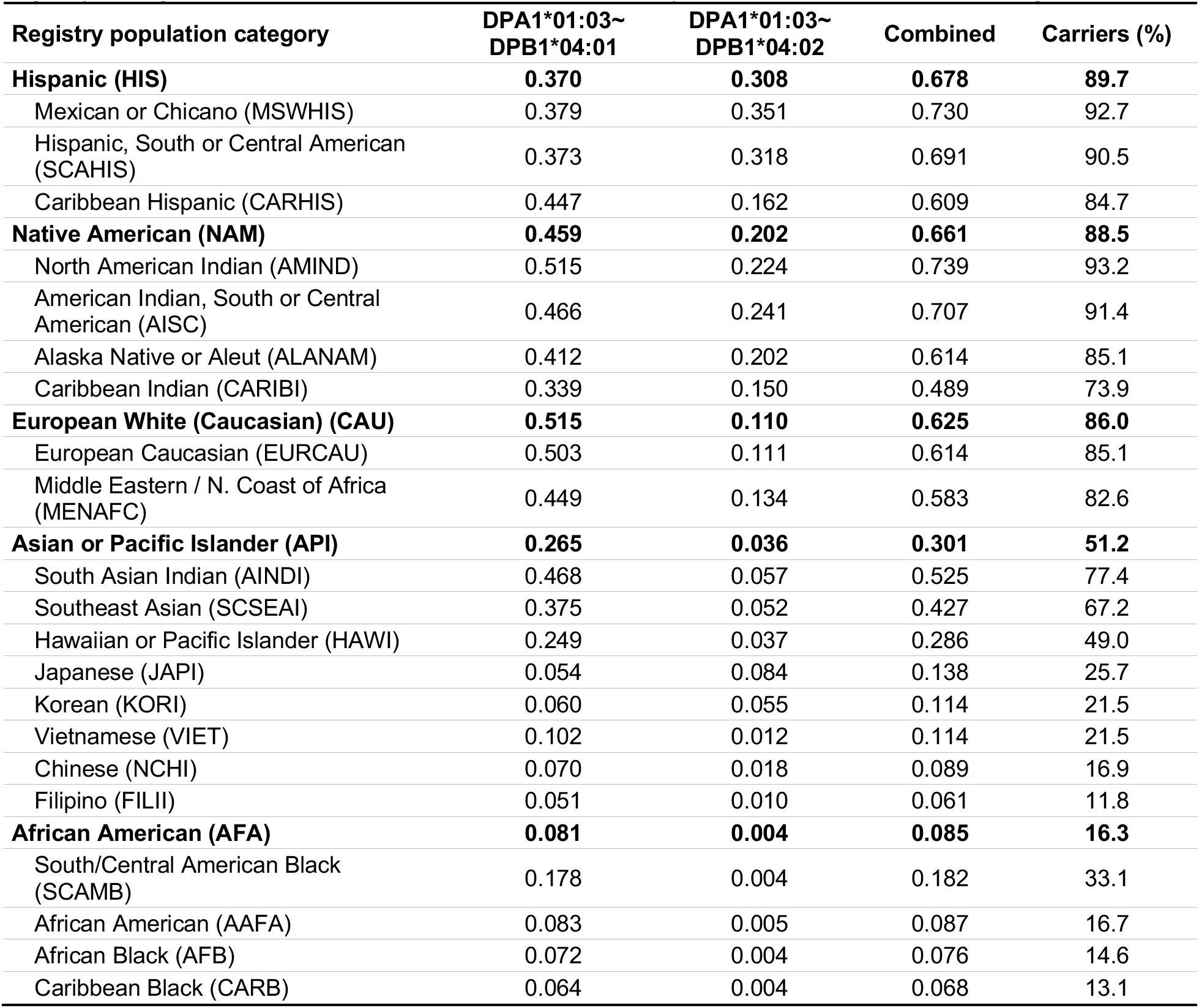
Estimated population prevalence in the United States of the HLA-DP04 haplotypes recognized by the NY-ESO-specific TCR. Frequencies of the DPA1*01:03/DPB1*04:01 and DPA1*01:03/DPB1*04:02 haplotypes in each registry population category, and the fraction of individuals expected to carry at least one of the two. Carrier frequency is 1 − (1 − f)², where f is the summed frequency of the two haplotypes, assuming Hardy–Weinberg equilibrium. Broad registry categories are shown in bold, each followed by its constituent detailed categories.

